# A comprehensive genome-scale model for *Rhodosporidium toruloides* IFO0880 accounting for functional genomics and phenotypic data

**DOI:** 10.1101/594846

**Authors:** Hoang V. Dinh, Patrick F. Suthers, Siu Hung Joshua Chan, Yihui Shen, Tianxia Xiao, Anshu Deewan, Sujit Sadashiv Jagtap, Joshua D. Rabinowitz, Christopher V. Rao, Huimin Zhao, Costas D. Maranas

## Abstract

**Background:** *Rhodosporidium toruloides* is a basidiomycetes yeast that can accumulate large amount of lipids and natively produce carotenoids. To better assess this non-model yeast’s metabolic capabilities, we reconstruct a genome-scale model of *R. toruloides* IFO0880’s metabolic network (*iRhto*1108) using recent functional genomics and phenotypic data in literature or generated herein.

**Results:** The model *iRhto*1108 accounts for 2,203 reactions, 1,985 metabolites and 1,108 genes. In this work, we integrate and supplement the current knowledge with in-house generated biomass composition and experimental measurements pertaining to the organism’s metabolic capabilities. Phenotype-genotype relationship predictions were improved through manual curation of gene-protein-reaction rules for 543 reactions and validations with gene essentiality data leading to correct recapitulations of 84.5% of gene essentiality data (sensitivity of 94.3% and specificity of 53.8%). Organism-specific macromolecular composition and ATP maintenance requirements were experimentally measured for two separate growth conditions: (i) carbon and (ii) nitrogen limitations. Overall, *iRhto*1108 reproduced *R. toruloides*’s utilization capabilities for 18 alternate substrates, matched measured wild-type growth yield, and recapitulated the viability of 772 out of 819 deletion mutants. As a demonstration to the model’s fidelity in guiding engineering interventions, the OptForce procedure was applied on *iRhto*1108 for the overproduction of triacylglycerol. Suggested interventions recapitulated many of the previously successfully implemented genetic modifications and put forth a few new ones.

**Conclusion:** *iRhto*1108 offers a highly curated model for a non-model yeast supported by multiple layers of experimental data that can be used to inform genetic interventions.

## Background

*Rhodotorula* genus species are found in various habitats including soil, water, air, on animals and plants, and even in extreme environments such as arctic ice sheets [1]. Among them, *Rhodosporidium toruloides* (or *Rhodotorula toruloides*) is a basidiomycete yeast generally found in soil [2] containing carotenoid compounds giving the organism its characteristic pink color [3, 4]. *R. toruloides* is an attractive metabolic engineering host for producing lipid and fatty acid-derived products due to its ability to accumulate lipid (predominantly triacylglycerols [5]) as high as 76% of the cell dry weight [6] and maintain lipid production in biomass hydrolysate containing growth inhibitory compounds [7]. It can also grow in high density cell cultures [8] and utilize a wide variety of substrates and glucose-xylose mixtures without catabolic repression [9, 10]. It has a rather compact genome (i.e., haploid genome of 20 Mb with 20% being intergenic sequence) that is tractable for genetic interventions [11–13]. Extensive metabolic engineering efforts have been focused on lipid production in *R. toruloides* by exploiting the organism’s ability to accumulate lipid under NaCl-enriched glucose-based media [14], nitrogen-limitation [15], sulfur-limitation [16], and phosphate-limitation conditions [17]. *R. toruloides* can accumulate lipids utilizing multiple substrates [9] and substrate mixtures [18]. Genetic interventions aimed at enhancing lipid accumulation have also been explored [11, 19] to overproduce fatty acid derived compounds (e.g., fatty alcohols and esters), used in surfactants, paints, and cosmetics [20, 21]. In addition to lipids, *R. toruloides* has also been used as a host for carotenoid [22] and d-arabitol production [23].

*R. toruloides* has recently been the target of significant research effort including genome (re)sequencing [3, 11, 13], functional genomics analyses [13], differential ‘omics characterization [3], determination of macromolecular composition [15], and growth kinetics in a continuous culture [24]. Collectively, these experiments have ushered an improved understanding of *R. toruloides* metabolism and provided the basis for the reconstruction of a metabolic model with genome-wide coverage. A comprehensive genome-scale metabolic reconstruction for *R. toruloides* would facilitate the integration of various heterogeneous datasets [25, 26] in making predictions of cellular phenotypes under various environmental and genetic perturbations and model-driven knowledge discovery [26], exploration of organism production potential [27–29], and extensions towards kinetic descriptions of metabolism [30, 31]. A successively improving sequence of metabolic models for *S. cerevisiae* have ushered significant insight into the organism’s physiology and offered many clues for re-engineering [32, 33]. Metabolic reconstruction of non-model yeasts have recently received significant attention, starting with *Pichia pastoris* for its use in production of recombinant protein [32] and the model oleaginous yeast *Yarrowia lipolytica* [20, 34] for which five genome-scale models of iteratively higher level of detail have been reconstructed [32, 35]. They were used to suggest fed-batch strategies to improve lipid accumulation and elucidate the regulation mechanism of lipid accumulation [36, 37]. We anticipate that similar advances would be spearheaded for *R. toruloides* facilitated by the genome-scale model described herein. In addition, to the benefits for guiding re-engineering efforts, a genome-scale model for *R. toruloides* will fill in a significant knowledge gap as the *Basidiomycota* phylum is highly under-represented in terms of metabolic model reconstructions. As of today, only a small metabolic model containing 85 reactions (without gene associations) [10, 18], and a genome-scale model associating with 897 genes [38] exist. In contrast, there exist numerous genome-scale models [32, 39] for eight organisms in the closely related *Ascomycota* phylum.

We hereby introduce the comprehensive genome-scale metabolic model of *R. toruloides* strain IFO0880, referred to hereafter as *iRhto*1108. It contains 2,203 reactions, 1,985 metabolites and spans 1,108 genes from the latest version of the genome [13]. The strain IFO0880 has been shown to be a robust host for lipid overproduction [11]. We used the model yeast 7.6 for *S. cerevisiae* [40] as the reconstruction process’ starting point and refined the draft reconstruction of the model by incorporating biochemical information from the latest genome annotation [13], biochemical (KEGG) information and the KBase database [41, 42]. By drawing from the gene essentiality results from a genome-wide functional genomic study [13], *iRhto*1108’s recapitulates 84.5% of the gene essentiality data. It contains highly curated gene-to-protein (GPR) associations including updates for 543 reactions (involving 373 genes) and confirmations of GPR assignments from *S. cerevisiae*. Using the conventions from GrowMatch procedure for benchmarking gene essentiality prediction [43], growth (G) and non-growth (NG) agreements or disagreements between model predictions and gene essentiality data are classified into four groups: G-G, G-NG, NG-G, and NG-NG; the first entry refers to the *in silico* result while the second part is the *in vivo*. The model achieved a sensitivity level of 94.3% (G-G / (G-G + NG-G)), a specificity level of 53.8% (NG-NG / (NG-NG + G-NG)), and an accuracy level of 84.5% ((G-G + NG-NG) / Total). Quantitative model predictions are also enhanced by organism-derived biomass compositions determined in this study under both carbon and nitrogen limited conditions revealing a much higher proportion of lipids in *R. toruloides* biomass compared to *S. cerevisiae*. ATP maintenance requirements were derived from a study on *R. toruloides*’s growth kinetics [24]. Metabolites present in the biomass reaction were extracted from both the original biomass equation of yeast 7.6, biomass constituent compendium [44] and the literature on *S. cerevisiae* biomass (see Methods). This compilation results in the addition of three cell wall components, nine cofactor and prosthetic groups, and seven metal ions. *iRhto*1108 contains 328 unique metabolic reactions (out of a total of 1,398 reactions) compared to both yeast 7.6 and the recent genome-scale model for *R. toruloides* strain NP11 version 1.1.0 [38]. The model predicts growth on all thirteen carbon substrates and five amino acids (as nitrogen source) that have been experimentally confirmed (see Supplementary Materials 2). Under nutrient starvation, *iRhto*1108 successfully captured *R. toruloides*’s lipid accumulation phenotype. As a demonstration of *iRhto*1108’s appropriateness to guide strain design, the OptForce procedure [45] was used on the model to pinpoint genetic interventions that led to an triacylglycerol overproducing phenotypes. Strain design solutions were in line with *in vivo* implemented flux “push-pull” strategy [46] that increased lipid production by approximately two-fold in *R. toruloides* [11]. Overall, *iRhto*1108 has undergone a detailed range of testing and validation studies promising to aid in future investigations of *R. toruloides*.

## Results and Discussion

### Model attributes and refinement of draft reconstruction

*iRhto*1108 is a comprehensive genome-scale model that integrates yeast biochemistry information from (i) previously built genome-scale models (*S. cerevisiae* yeast 7.6 [40], (ii) KBase fungal models [42]), and (iii) *R. toruloides* specific information extracted from the primary literature [13, 23, 47] or generated herein. The model statistics are summarized in Table 1. Genes included in *iRhto*1108 cover 13% of the organism’s chromosomal and 6% of mitochondrial genome. Throughout this article, genes named will be referred to by the corresponding *S. cerevisiae*’s homolog name (if available) (e.g., *HOM6* for homoserine dehydrogenase) or otherwise using *R. toruloides* gene IDs (e.g., *rt6880* for serine O-acetyltransferase). *R. toruloides* protein IDs are not used herein (e.g., *RTO4_15248* for serine O-acetyltransferase) to retain consistency in gene identification. *iRhto*1108 shares 65% of genes, 70% of reactions, and 67% of metabolites with the *cerevisiae* yeast 7.6 model [40]. KBase entries contributed 8% of genes, 5% of reactions, and 7% of metabolites of *iRhto*1108. KBase was used to identify additional homologous genes and extract reactions from metabolic reconstructions for non-model yeasts. The remainder of the model content (i.e., 27% of genes, 25% of reactions, and 26% of metabolites) was directly culled from the genome annotation and subsequently manually curated. Many of these model additions do not necessarily capture *R. toruloides*-only biochemistry but instead unpack aggregated yeast 7.6’s reaction content or replace redundant yeast 7.6’s features. For example, using KBase, a lumped palmitoyl-CoA synthesis (fatty acid C16:0) reaction in yeast 7.6 is detailed into 28 steps catalyzed by fatty acid synthase in *iRhto*1108 (seven elongation cycles, each cycle contains four elementary steps). In addition, a set of aggregated reactions simplified lipid metabolism in yeast 7.6 [40, 48]. For example, a single generalized reaction for diacylglycerol acyltransferase (DGAT_rm) replaced 32 copies of DGAT_rm in yeast 7.6 operating on 32 variants of triacylglycerol. Overall, gene-protein-reaction associations were assigned for 93% of metabolic reactions in *iRhto*1108. Not surprisingly, identification of genes coding for transporters remained a challenge as was the case for yeast 7.6. Missing GPR assignments in *iRhto*1108 are mostly in intracellular transport between compartments (i.e., 364 from a total of 456 GPR-lacking reactions). Throughout the reconstruction process, we manually curated the GPR of 543 reactions associated with 373 genes. Determination of metabolic role of a gene(s) in GPRs was assisted using NCBI’s Conserved Domain Database (NCBI’s CDD) [49]. For example, initially no reaction enabling the synthesis of spermidine, a biomass constituent, was found as no spermidine synthase gene was identified via bidirectional BLAST. However, subsequent analysis by NCBI’s CDD revealed a catalytic domain on *rt8465* (homolog of *S. cerevisiae*’s *LYS9*). Notably, this catalytic domain does not overlap with saccharopine dehydrogenase domain identified by bidirectional BLAST. Thus, the spermidine synthase reaction was subsequently determined to associate to *rt8465* and thereby fill the gap in spermidine synthesis. Furthermore, using the more recent version of the consensus yeast model available at (https://github.com/SysBioChalmers/yeast-GEM, yeast 8.3.3), GPR assignments for 22 reactions in *iRhto*1108 were updated. For example, drawn from yeast 8.3.3’s GPR assignment, isozyme *rt1542* (homologous to *YOR283W*) was added to the glycolysis reaction phosphoglycerate mutase’s GPR.

**Table 1.** *R. toruloides iRhto*1108 genome-scale model statistics

| Properties | Statistics |
| --- | --- |
| **Genes** | 1,108 |
| Identified from yeast 7.6 | 717 |
| Identified from KBase | 86 |
| From chromosome (% of chromosomal ORFs) | 1,087 (13%) |
| with *S. cerevisiae* homolog identified | 908 |
| From mitochondrial genome (% of mitochondrial ORFs) | 21 (6%) |
| with *S. cerevisiae* homolog identified | 7 |
| **Reactions** | 2,203 |
| From yeast 7.6 | 1,514 |
| From KBase | 118 |
| Metabolic reactions | 1,398 |
| With GPR assigned | 1,306 |
| Unique number of metabolic reactions | 1,129 |
| Transport reactions | 619 |
| Extracellular transport | 163 |
| with GPR assigned | 75 |
| Intracellular transport | 456 |
| with GPR assigned | 92 |
| Exchange reactions | 185 |
| **Metabolites** | 1,985 |
| From yeast 7.6 | 1,328 |
| From KBase | 145 |
| Unique metabolites | 1,043 |
| Formula and charge assignments from database | 1,535 |
| **Compartments** | 14 |

The classification of *iRhto*1108 genes shown in Table 1 quantifies the extent of contribution from previous yeast reconstructions and the significant expansion in *iRhto*1108 (Figure 1). Eukaryotic orthologous group (KOG) assignments [50], a Eukaryotic-specific Cluster of orthologous groups of protein (COG) provided by the updated genome annotations [13], were used in classifying genes and the associated reactions. Most of the genes in *iRhto*1108 are homologous to *S. cerevisiae*’s genes (Figure 1), in agreement with the relative phylogenetic proximity between the two species of yeasts (i.e., their respective divisions, Ascomycota and Basidiomycota, are grouped to the sub-kingdom Dikarya). Highly conserved metabolic functions (i.e., 80-98% genes with homologs identified per KOG class) between the two are observed in all the core metabolic functions (listed in decreasing degree of conservation): nucleotide, inorganic ion, cell wall, coenzyme, amino acid, carbohydrate, energy production, and lipid metabolism. Nevertheless, *R. toruloides* has a number of unique metabolic capabilities compared to *S. cerevisiae*. These new functionalities, 328 metabolic reactions out of 1,398, are not predominantly localized in any specific pathway but rather span multiple KOG classifications. In terms of genome coverage, *iRhto*1108 is able to account for 66-77% of genes in KOG classes, namely energy nucleotide, coenzyme, amino acid, and energy production metabolism. The category with the lowest genome coverage (i.e., 25%) is inorganic ion transport and metabolism. A significant fraction of *iRhto*1108’s genes fell into the “Unassigned” category due to KOG’s inability to identify genes with non-homologous sequences that perform core metabolic functions. For example, *R. toruloides* fatty acid synthase subunit I and II were not classified by KOG annotation into the lipid metabolism group, possibly due to irregular arrangement of catalytic sequence motifs compared to *S. cerevisiae* and other types of yeast [6].

**Figure 1.**
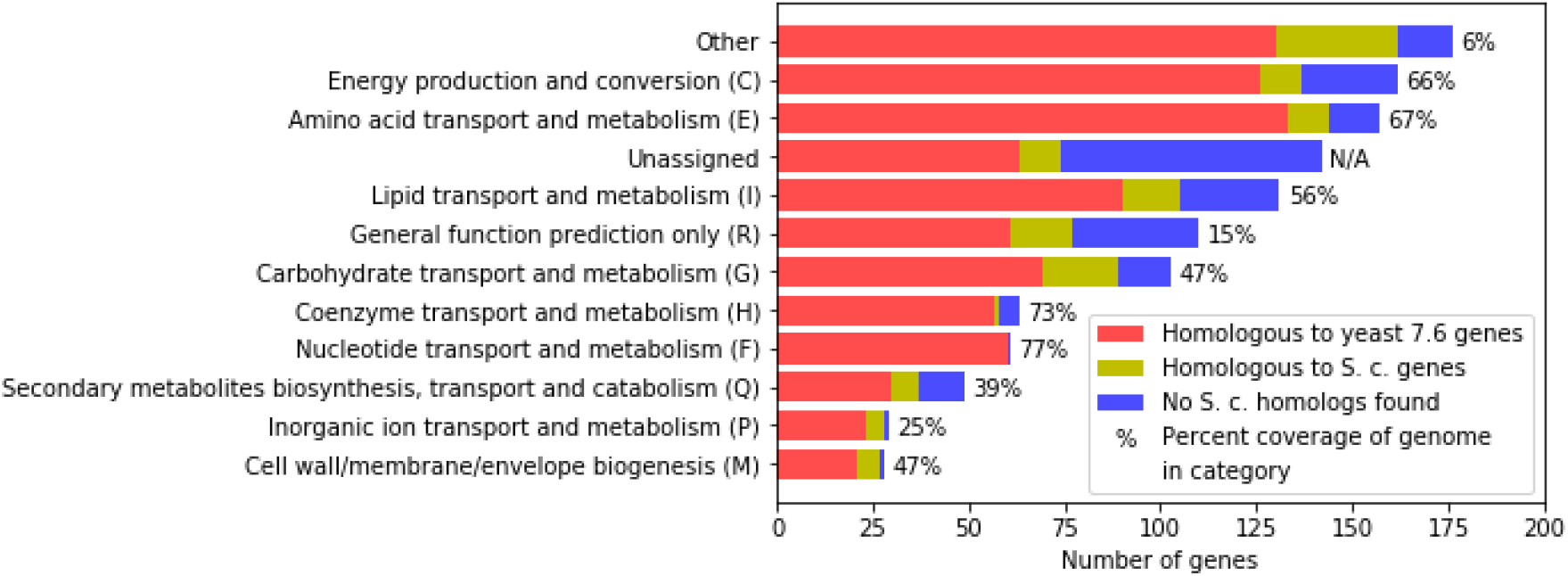
Classifications of genes in *iRhto*1108. Eukaryotic orthologous groups (KOG) annotations are provided in the genome annotation and used for classifying genes to the corresponding functions. Group abbreviations are in the parentheses. A gene with multiple KOG groups assignments were added to all the groups. A gene without KOG annotation was manually assigned to a KOG group. Other groups include B, D, J, K, L, N, O, S, T, U, V, Z, W, Y, and A (see https://genome.jgi.doe.gov/Tutorial/tutorial/kog.html).

The majority of the novel metabolic functions captured in *iRhto*1108 are extracted directly from the genome annotation or open literature. For example, included in the model are reactions and associated genes for both the carotenoid [47] and d-arabitol production pathways [23] which were absent in yeast 7.6. Most of the novel functions belong to lipid and carotenoid metabolism. Some examples in lipid metabolism are ATP citrate lyase in acetyl-CoA production [51], cytoplasmic malic enzyme [11], and stearoyl-CoA desaturase in polyunsaturated acyl-CoA production [19]. The carotenoid biosynthesis pathway whose products are responsible for the organism’s characteristic pink color [4] is captured in *iRhto*1108. In addition, an NADH oxidoreductase reaction (complex I in electron transport chain) is included in *iRhto*1108 which is known to be absent in *S. cerevisiae* [52]. Non-essential metabolic functions found in the genome annotation with an unclear physiological role such as a peroxisomal d-amino acid oxidase [53] are also recorded in the model.

Model annotation and network consistency are important properties for testing genome-scale model quality [54]. We evaluated *iRhto*1108 using standardized tests provided by the memote test suite and updated the model based on detected issues. Under the independent section (scored tests), *iRhto*1108 received a high score of 87% on biochemical annotation and network consistency tests. Since *R. toruloides* genome is recorded on JGI Mycocosm and not on memote-verified databases such as KEGG, we only provide the *S. cerevisiae* homologs as genes’ annotation. *iRhto*1108 also achieve a score of 99.7% on network consistency tests. Some lost points are from memote’s mistaken identification of reaction imbalances, namely generalized reactions that set the composition of generic acyl-CoA (using “Acyl” as a group in the formula) and biochemical reactions associated with that generic acyl-CoA group. Detected by memote, unbounded fluxes that can form a thermodynamically infeasible cycle [55] were fixed by restricting the directionality of transporters and/or reactions. For example, the tyrosine importer (*AVT1*), exporter (*AVT3*), and efflux transporter (*ATG22*) from cytosol to vacuole can shuttle tyrosine in and out of the vacuole with no driving force leading to an unbounded flux. To remedy this, the efflux transporter was allowed to only import (not export) tyrosine to the vacuole. Export function can be re-activated if needed (e.g., under autophagy-induced protein degradation [56]). In addition, under network topology tests, memote reported a high number of blocked reactions (677 out of 2,203). These reactions and metabolites were retained in *iRhto*1108 as they are based on assignments from homologous genes and genome annotation. They cause no problems in flux balance analyses and may serve in the future as gapfilling targets.

### Update of biomass composition and revision of ATP maintenance requirements

In addition to differences in pathways, an important contribution in *iRhto*1108 is the expansion of the list of biomass constituents by 23 components from the original 45 taken from yeast 7.6 (Table 2). Four new fatty acid species, C18:2, C18:3, C20:0, and C24:0, are added to the biomass reaction based on their detection by mass spectrometry measurements (see Methods). Seven metal ions are added based on biomass measurements for *S. cerevisiae* [57]. We identified three cell wall components and nine cofactors and prosthetic groups that must be added to match measured phenotypes [13] (see Methods). For instance, lethal knockouts of dephospho-CoA kinase (gene: *CAB5*) or GPI anchor biosynthesis (gene: *GPI13*) (KEGG reaction R05923) are unresolvable without the additions of coenzyme-A and GPI anchor to the biomass reaction, respectively. These validations are provided in Supplementary Materials 2. The revised list of biomass constituents and the experimentally determined macromolecular composition are provided in Table 2. The full description of the biomass reaction is detailed in the Supplementary Materials 1. Moreover, we experimentally determined *R. toruloides* macromolecular composition separately under both carbon and nitrogen limitation (see Table 2 and Methods section). Protein, carbohydrate, DNA, RNA, and lipid composition were measured for cells growing on glucose in a chemostat at the dilution rate of 0.1 hr^−1^. Further improvements include DNA base composition update based on GC content [58], RNA bases composition informed from RNA-Seq data, and lipid’s acyl group composition measured by mass spectrometry (see Methods). Both biomass reactions for *R. toruloides* in carbon and nitrogen limited conditions imply a higher proportion for lipid than *S. cerevisiae* (in comparison to yeast 7.6’s biomass reaction) and a lower proportion of carbohydrate (and protein under carbon limitation). The DNA fraction for *R. toruloides* is also higher under both conditions compared to *S. cerevisiae*. Both the lipid and DNA fractions are higher while the RNA fraction is lower for cells growing under nitrogen limitation. Importantly, the combined coefficient-weighted molecular weights of all constituents were standardized to 1 g.mmol^−1^ to ensure consistency of growth yield prediction [59]. The biomass composition listed in Table 2, follows the core biomass definition [60, 61] consisting of growth-essential metabolite requirements. The inclusion of these metabolites in the biomass reaction was based on gene essentiality results [13] and experimental data from *S. cerevisiae* (Supplementary Materials 2). Compared to rhto-GEM model v. 1.1.0 [38], *iRhto*1108’s biomass reaction contains 23 additional constituents. In addition, *iRhto*1108 offers a nitrogen limited version (viz., conditions applicable for lipid production) underpinned by a significant biomass compositional difference.

**Table 2.** Summary of *iRhto*1108’s biomass composition.

| Constituents |  |  | Composition (%) |  |  |
| --- | --- | --- | --- | --- | --- |
|  |  |  | C-lim | N-lim | yeast7.6 |
| *Protein*<sup>a</sup> |  |  | 43.31 | 30.71 | 35.71 |
| L-Alanine | L-Arginine | L-Asparagine |  |  |  |
| L-Aspartate | L-Cysteine | L-Glutamine |  |  |  |
| L-Glutamate | Glycine | L-Histidine |  |  |  |
| L-Isoleucine | L-Leucine | L-Lysine |  |  |  |
| L-Methionine<br>L-Serine<br>L-Tyrosine | L-Phenylalanine<br>L-Threonine<br>L-Valine | L-Proline<br>L-Tryptophan |  |  |  |
| *Carbohydrate*<br>1,3-beta-D-Glucan<br>**N-Glycan<sup>b</sup>** | 1,6-beta-D-Glucan<br>**O-Glycan<sup>b</sup>** | Chitin<br>**GPI-anchor<sup>b</sup>** | 32.61 | 12.50 | 52.27 |
| *Lipid*<br>Episterol<br>Phosphatidylcholine<br>Phosphatidylserine | Free fatty acids (7 species)<sup>c</sup><br>Phosphatidylethanolamine<br>TAG | Inositol-P-ceramide<br>Phosphatidylinositol | 12.33 | 44.84 | 0.74 |
| *RNA<sup>d</sup>*<br>ATP<br>UTP | CTP | GTP | 6.73 | 4.69 | 5.85 |
| *DNA<sup>d</sup>*<br>dATP<br>dTTP | dCTP | dGTP | 1.12 | 3.36 | 0.34 |
| *Cofactors and prosthetic groups*<br>**S-Adenosyl-L-methionine**<br>**FAD**<br>**NADP**<br>**Tetrahydrofolate** | **Biotin**<br>Heme A<br>Riboflavin<br>**Thiamine diphosphate** | **Coenzyme-A**<br>**NAD**<br>**Spermidine** | 0.06 | 0.06 | 0.03 |
| *Inorganic ions*<br>**Calcium**<br>**Magnesium**<br>**Potassium** | **Copper**<br>**Manganese**<br>Sulphate | **Iron**<br>Phosphate<br>**Zinc** | 3.85 | 3.85 | 5.06 |

Biomass constituents absent from yeast 7.6 are shown in boldface type. Different representations of yeast 7.6 biomass constituents are listed in the notes below. A detailed analysis of *iRhto1108*'s biomass composition is provided in the Methods section and the full description in the Supplementary Materials 1.
<sup>a</sup> Identical to those in yeast 7.6, amino acids in the biomass objective function are in charged-tRNA form.
<sup>b</sup> The generic mannan (mannose-containing) metabolite in yeast 7.6 was replaced with three specific essential cell wall components [63].
<sup>c</sup> Seven free fatty acid species were abundant (>1% weight) in growth experiments detailed in the Methods section. These are palmitate (C16:0), stearate (C18:0), oleate (C18:1), linoleate (C18:2), linolenate (C18:3), docosanoate (C22:0), and tetracosanoate (C24:0).
<sup>d</sup> Monophosphate ribonucleic and deoxyribonucleic acids in yeast 7.6 were replaced with the corresponding triphosphate ones.

In addition to the updated biomass composition derived for two separate growth conditions, we also revisited the ATP maintenance requirements (both growth and non-growth). Non-growth (NGAM) and growth associated maintenance (GAM) values were estimated by assessing the model’s optimal ATP production under glucose uptake restriction and growth yield requirement and experimentally recorded glucose uptake rates and growth rates. Correctly assessing ATP maintenance is important for properly quantifying energetic needs and growth yield [61]. ATP maintenance requirements for *iRhto*1108 were calculated from available chemostat data for growth on glucose for both carbon and nitrogen limitation, respectively [24] (see Methods). An NGAM value of 1.01 mmol gDW^−1^ hr^−1^ for both conditions was recovered. In contrast, the growth associated maintenance (GAM) was condition-dependent with a value of 140.98 mmol gDW^−1^ under carbon limited and 276.37 mmol gDW^−1^ under nitrogen limited conditions. In yeast 7.6, NGAM is not modeled (though an earlier *S. cerevisiae* model [62] reported an NGAM value of 1 mmol gDW^−1^) and the GAM value is 59.28 mmol gDW^−1^. The GAM value quantifies growth-associated energy costs that are not captured in the biomass equation, alluding to higher energy demands for *R. toruloides* growth compared to *S. cerevisiae*. Under nitrogen limitation, GAM is even higher (1.9-fold increase). Note that growth kinetics of *R. toruloides* under nitrogen limitation follows a different trend compared to under carbon limitation [24]. It appears that the assumption of constant GAM value across all growth rates may not hold under nitrogen limitation. However, higher ATP cost under nitrogen limitation is generally accepted (see Supplementary Materials 2). In rhto-GEM model v. 1.1.0 [38], a non-condition-specific GAM value of 131 mmol gDW^−1^ and NGAM value of 3 mmol gDW^−1^ hr^−1^ were reported. These values generally match the *iRhto*1108’s corresponding entries under carbon limitation.

### Gene essentiality, growth viability, and phenotype predictions

*iRhto*1108 predictions were contrasted against gene essentiality and mutant auxotrophy data derived from the functional genomics study [13] and growth yield and viability data from multiple literature sources (see Supplementary Materials 2). The data collected and corresponding predictions are summarized in Table 3. Gene sequence disruptions with T-DNA insertions were carried out leading to evaluation of gene essentiality for 1,079 of the 1,108 genes in the model [13]. Gene essentiality predictions are shown in Table 3. *iRhto*1108 achieved 84.5% accuracy (i.e., correct prediction of gene essentiality and non-essentiality over all predictions) for gene essentiality prediction which is similar to that of yeast 7.6 (i.e., 89.8%). The model is particularly adept at recovering mutant growth, measured by the sensitivity level of 94.3% Positive mutant growth misses (NG-G) (i.e., 5.5%) by *iRhto*1108 were mainly due to differences in protein subunit assignments between *S. cerevisiae* and *R. toruloides*. For example, consistent with *S. cerevisiae* but in contrast to *R. toruloides*, *GPI2* and *GPI15* knockouts were predicted by *iRhto*1108 to be lethal since they are subunits of the enzyme catalyzing the first step of GPI anchor biosynthesis. The resolution of this mismatch would require customizing the GPR relationship for *R. toluloides* instead of simply carrying the one from *S. cerevisiae*. *iRhto*1108 relatively inaccurate predictions of negative mutant growth (G-NG) (i.e., specificity of 53.8%) matches the corresponding specificity for yeast 7.6 (i.e., 52.5%) as the GPR assignments were directly ported from yeast 7.6. For example, for L-methionine auxotrophic (i.e., *rt6880* and *rt3663*) mutants upon knocking out cysteine biosynthesis via serine [13], *iRhto*1108 predicted growth without the need for L-methionine supplementation by allowing sulfur assimilation via the homoserine pathway. The lack of observed growth suggests the possible inactivity of the homoserine pathway (encoded by *MET2*, *6*, and *17*).

**Table 3.** Summary of phenotype predictions by *iRhto*1108.

| Prediction | Statistics |
| --- | --- |
| **Gene essentiality<sup>a</sup>** |  |
| G-G | 772 (72%) |
| G-NG | 120 (11%) |
| NG-G | 47 (4%) |
| NG-NG | 140 (13%) |
| Accuracy | 84.5% |
| Sensitivity | 94.3% |
| Specificity | 53.8% |
| **Growth viability** |  |
| As carbon source | 13 |
| As nitrogen source (positive) | 5 |
| As nitrogen source (negative) | 1 |
| **Mutant auxotrophy** |  |
| Arginine and/or methionine | 18 / 22 |
| **Growth yield prediction** |  |
| D-Glucose | 5 validations |
| D-Xylose | 9 validations |
| **Production yield prediction** |  |
| D-Arabitol | 6 validations |

<sup>a</sup> Agreements or disagreements between model predictions and gene essentiality data are classified into four groups: G-G, G-NG, NG-G, and NG-NG; the first part of the group is in silico result, the second part is in vivo result, “G” stands for growth, and “NG” stands for non-growth. Accuracy = (G-G + NG-NG) / Total. Sensitivity = G-G / (G-G + NG-G). Specificity = NG-NG / (NG-NG + G-NG).

*iRhto*1108 was also tested in terms of its ability to predict growth on alternative substrates (see Table 3). The simulations were performed using the model with substrate uptake rate and secretion rate as inputs. *R. toruloides* can grow on d-glucose, d-xylose, acetate, glycerol, fructose, mannose, sucrose, cellobiose and fatty acids as carbon sources. Amino acids such as L-threonine, L-serine, L-proline, L-alanine, and L-arginine can support *R. toluloides* nitrogen needs. Growth experiments on cellobiose, mannose, sucrose, and amino acids were performed in this study whereas the other findings are collected from the literature (see Supplementary Materials 2). *iRhto1108* predicts growth with these substrates upon activating the corresponding transporters. The activation of L-threonine transporter is seemingly in conflict with gene essentiality results since knockouts in *de novo* L-threonine biosynthesis are lethal in rich media (five steps from L-aspartate each encoded by *HOM3*, *HOM2*, *THR1*, *THR4*, and *HOM6*) [13]. Further study on L-threonine biosynthesis pathway is necessary to explain why *ex vivo* L-threonine supplementation was insufficient to rescue those mutants. Other growth phenotypes such as mutant auxotrophy and quantitative growth yields are collected and used in model validations. Arginine and methionine auxotrophy predictions for 22 mutants are largely consistent with experimental findings [13] and explanations for inconsistencies (4 mutants) can be offered based on the model predictions such as the hypothesis of homoserine pathway being repressed under L-methionine abundance conditions, as mentioned above. Furthermore, *iRhto*1108 quantitative prediction of wild-type growth yield in exponential phase are close to the experimental numbers (see Supplementary Materials 2).

### Phenotypic change under nutrient starvation conditions

Nutrient starvation is a common strategy for enhancing lipid accumulation in oleaginous yeasts. It has been successfully implemented in *R. toruloides* [15], *Y. lipolytica* [37], and *Lipomyces starkeyi* [64]. *R. toruloides* also over-accumulates lipid under phosphate [17] and sulphate [16] limitation. Triacylglycerol (TAG), the major compound in lipid accumulation, is stockpiled in lipid particles thus sequestering excess carbon substrate. We sought to recapitulate lipid accumulation in response to nutrient limitation conditions in *iRhto*1108 using two separate maximization problems. First, biomass production was prioritized by setting the flux balance analysis objective function to be maximization of growth yield given a nutrient-limited input. Second, lipid accumulation under low nitrogen was imposed to *iRhto*1108 by setting the model objective function to be maximization of TAG production. This optimization posture is hypothesized to be the regulatory outcome involving the TOR signaling pathway in *R. toruloides* NP11 [3]. Using these two maximization problems both growth and TAG yield were calculated under varying degrees of limitation for inorganic ions (i.e., ammonium, phosphate, and sulphate) and oxygen. Dissolved oxygen is an experimentally controllable variable that has been shown to affect lipid accumulation [65]. In an experiment in which ammonium was limited for *Y. lipolytica*, reduced aeration rate was shown to enhance lipid production [36]. Phenotype phase plane analysis [66] has been used to examine growth and TAG yield with respect to these two sources of variation.

Under nutrient limitation, *iRhto*1108 predicts an increase in TAG yield and decrease in biomass yield which is in qualitative agreement with experimental observations for *R. toruloides* and other oleaginous yeast. The same trend is observed under nitrogen, phosphate, and sulphate limitation This suggests that cell proliferation is the primary cellular objective for *R. toruloides* in the absence of nutrient limitation whereas TAG storage becomes the objective function for *iRhto*1108 in response to nutrient limitation conditions. The effect of oxygen limitation on TAG production in *iRhto*1108 follows a more complex trend depending on the degree of nutrient limitation. Moderate reduction in oxygen availability decreased growth yield with no effect on TAG yield. However, when oxygen level is reduced below a threshold (see the diagonal gradient region on Figure 2, column A), TAG production is also compromised and growth yield is directly proportional to oxygen level (Figure 2, column B). Note that reduction in TAG yield was also observed experimentally in *Y. lipolytica* under severe oxygen limitation with N_2_ aeration [36]. This is not surprising as oxygen is necessary to provide enough energy through oxidative phosphorylation for TAG synthesis. TAG biosynthesis requires one NAD(P)H for dihydroxyacetone reduction and an ATP plus two NADPH molecules for every two-carbon elongation step of the acyl groups. Acetyl-CoA biosynthesis required for fatty acid biosynthesis also consumes an ATP. Moderate reduction of oxygen availability did not improve TAG yield based on the phenotype phase plane analysis (Figure 2). The result is in agreement with experimental results for *R. toruloides* [18] but it is in contrast with results for *Y. lipolytica* [36]. Overall, TAG accumulation appears to be insensitive to moderate oxygen availability reduction but becomes highly bottlenecked under very low oxygen availability. In contrast, growth is affected by both oxygen and ammonium availability reaching a maximum when both are in excess.

**Figure 2.**
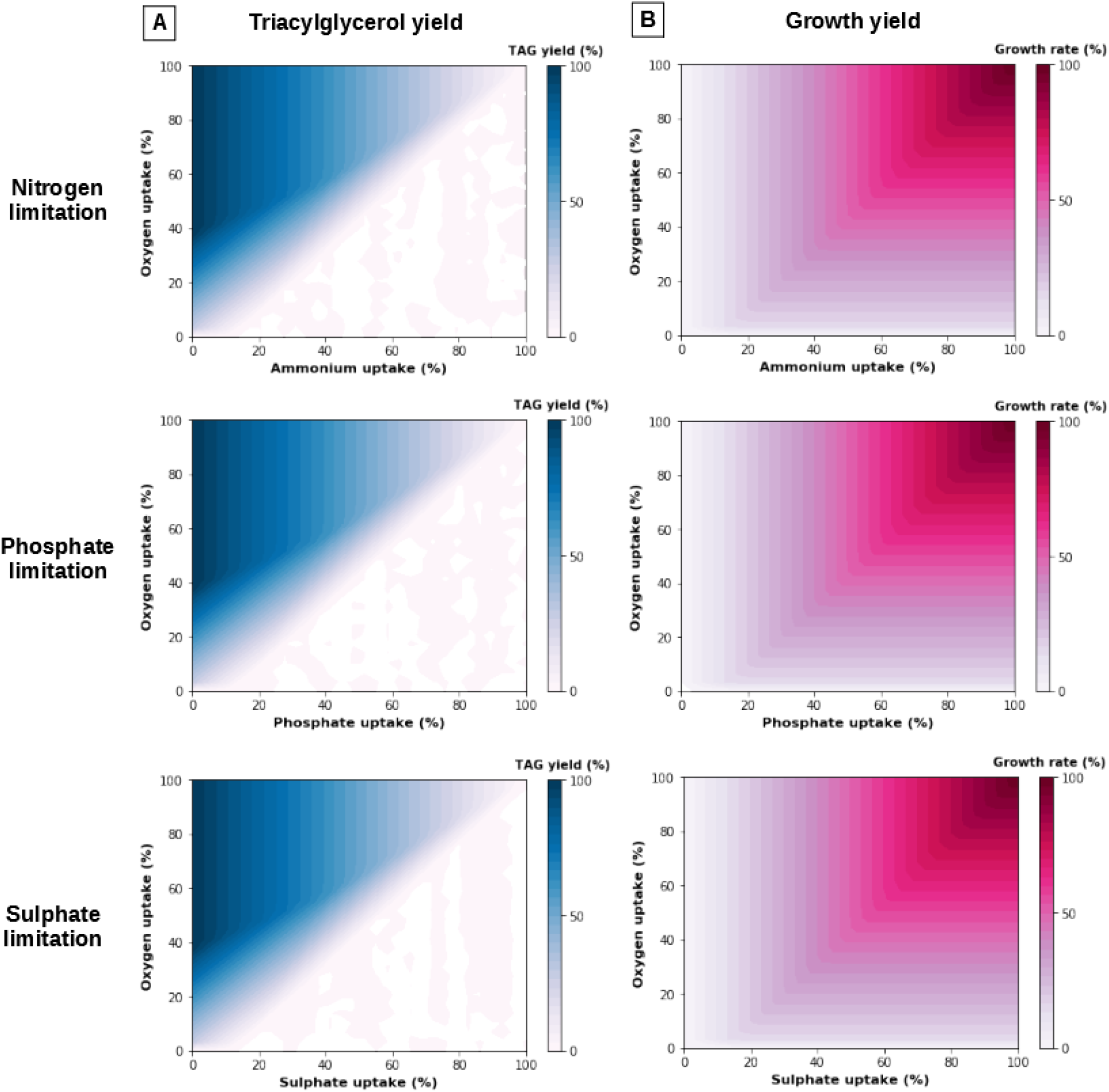
Phenotype phase planes of TAG production (column A) and maximal growth yield (column B) in nutrient (i.e., ammonium, phosphate, sulphate) and oxygen limited conditions. Values on the figure are percentage of maximal allowed flux for nutrients uptake and maximal yield for TAG production and growth rate. Determined by the model, upper bounds of uptake values are minimal amount required to sustain maximal growth (oxygen 12.78, ammonium 2.43, phosphate 0.20, and sulphate 0.03 mmol.gDW^−1^.hr^−1^). Maximal TAG production is 0.31 g/g glucose and maximal growth rate is 0.38 hr^−1^.

**Figure 3.**
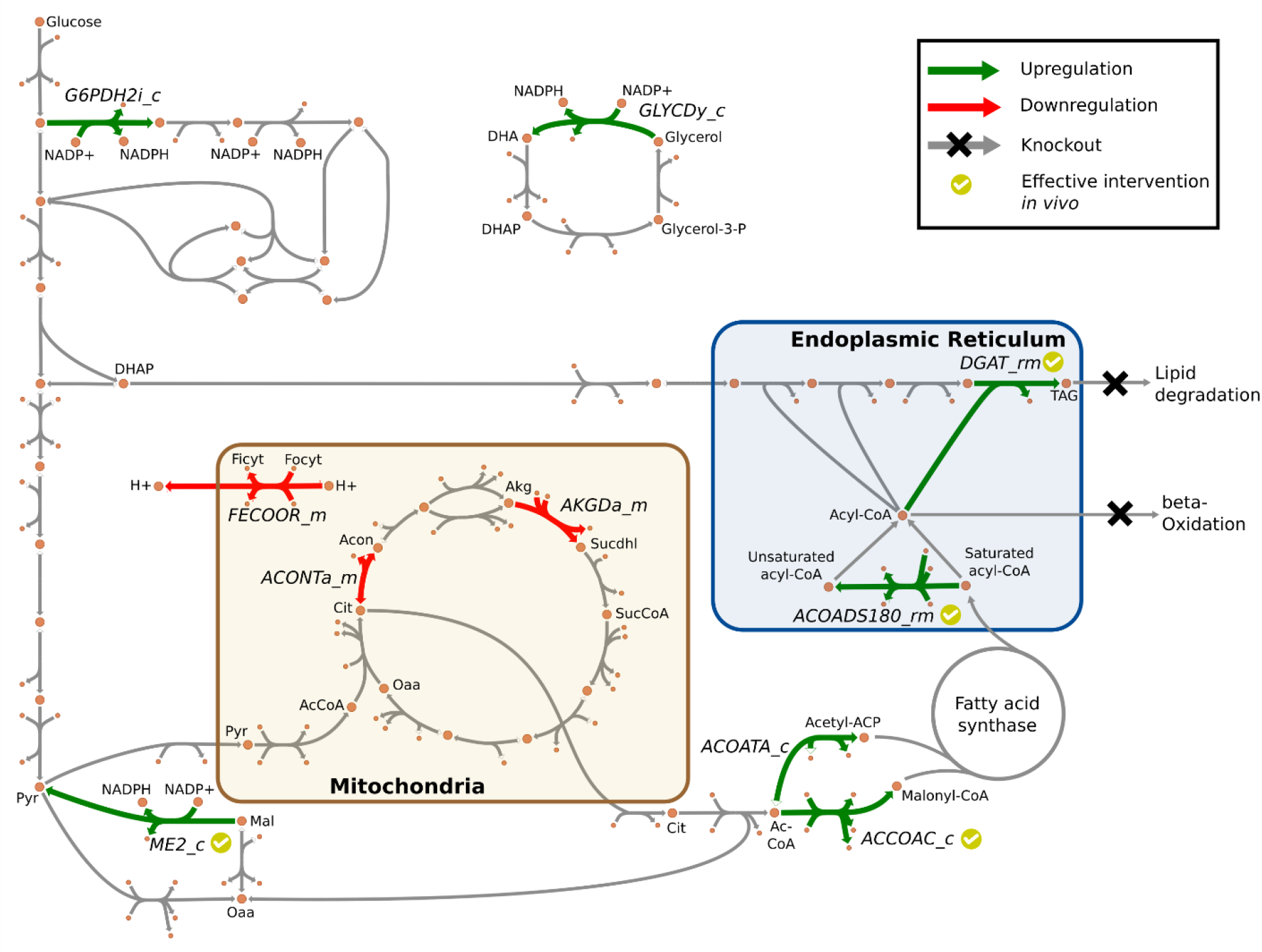
Visualization of triacylglycerol production pathway. Interventions identified by OptForce and implemented *in vivo* were annotated. Reaction abbreviations are listed in Table 4 and detailed in Supplementary Materials 1. Metabolite abbreviations: DHA – dihydroxyacetone, DHAP – DHA phosphate, Ficyt – ferricytochrome, Focyt – ferrocytochrome, Pyr – pyruvate, Mal – malate, AcCoa – acetyl-CoA, Oaa – oxaloacetate, Cit – citrate, Acon – aconitate, Akg – alpha-ketoglutarate, Sucdhl – S(8)-succinyldihydrolipoamide, SucCoA – succinyl-CoA, TAG – triacylglycerol.

As described above, *iRhto*1108 captures changes in phenotype for *R. toruloides* under nutrient starvation. We next explored whether the corresponding flux changes for a new phenotype quantitatively match gene upregulation/downregulation data [3] under the carbon and nitrogen limited version of the model (*iRhto*1108C and *iRhto*1108N, respectively). We used as a criterion of gene-reaction correlation that the change in gene expression level and associated reaction flux change is within a factor of two. In addition, genes with a p-value of less than 0.05 and a false discovery rate of less than 0.001 were excluded from the analysis as proposed in Zhu et al., 2012 [3]. We identified 12 upregulated and 11 downregulated genes under nitrogen starvation that quantitatively matched changes in metabolic fluxes predicted by *iRhto*1108. Overall, we find very few genes (23 out of 1,064) where the change in mRNA level quantitatively tracks the shift in model-predicted metabolic fluxes. Gene expression levels (taken from Zhu et al., 2012 [3]) and metabolic flux values are reported in Supplementary Materials 2. Increased energy demands under nitrogen limitation, was accompanied by upregulation for subunits of ATP synthase (*ATP14*, *16*, and *VMA9*), NADH:ubiquinone oxidoreductase (complex I) (*NDE2*, *rt0331*, *rt4846*, *rt1642*, and *rt2984*), and ferrocytochrome-c:oxygen oxidoreductase (complex III) (*rt2984*). In addition, increased lipid fraction of biomass under nitrogen limitation, is consistent with upregulation of genes in sterol (*ERG7*, *12*) and sphingolipid synthesis (*rt3023*). In contrast, a gene in phospholipid synthesis were downregulated (i.e., *CHO1*) which is consistent with the lower phospholipid fraction among lipid species under nitrogen limitation [15]. The reduced biomass fraction towards carbohydrate fraction was accompanied by downregulations in cell wall biosynthesis (*GSC2*, *ALG1*, *GFA1*, *GPI13*, *GPI14*, *SEC53*, *rt1388*, *CHS1/2*, and *MNT3*). Overall, only a few genes (23 out of 1,064) quantitatively tracked the corresponding flux changes. This re-emphasizes that flux distribution does not simply track changes in gene expression but rather are affected by many other factors such as transcriptional regulation, translation efficiency, substrate level regulation, metabolite pools, etc. [67].

### Predicting metabolic engineering strategies for enhanced triacylglycerol production

In this section, we explore the effectiveness of *iRhto*1108 to guide strain design by contrasting predictions for TAG overproduction with successful experimental interventions. A number of re-engineering strategies have been recently implemented in *R. toruloides* by mimicking effective interventions in *Y. lipolytica* [68]. These interventions include upregulation of acetyl-CoA carboxylase, diacylglycerol transferase, malic enzyme, and stearoyl-CoA desaturase [11, 19]. We used the OptForce procedure [45] using *iRhto*1108 as the metabolic map and contrasted with existing solutions. The goal was not necessarily to find new interventions but rather to assess whether *iRhto*1108 can indeed steer strain design algorithms towards promising designs. The OptForce procedure was applied for TAG overproduction using glucose as the carbon substrate (see Methods). ^13^C metabolic flux analysis (MFA) data for *Y. lipolytica* under nitrogen limitation [69] was used as a stand-in to determine the reference flux distribution for wild-type strain as no such data is currently available for *R. toluloides*. *Y. lipolytica* is a closely related oleaginous yeast with similar metabolic capability such as utilizing ATP citrate lyase for cytoplasmic acetyl-CoA production. Nitrogen availability level of 33% (of the maximum needed) was inferred from the MFA flux data by calculating the minimal amount of ammonium uptake. The model built for nitrogen limited conditions, *iRhto*1108N, was used throughout the OptForce simulation.

First, sets of candidates for overexpression (MUST^U^), downregulation (MUST^L^), and knockout (MUST^X^) were determined by contrasting the flux ranges of wild-type and overproducing strain. We excluded *in vivo* essential reactions [13] from MUST^X^ and transport, exchange, and generalized reactions from all MUST sets. For a sequence of reactions in series only the first step was considered as a perturbation candidate. For example, among 31 reactions of the fatty acid synthase’s chain elongation, only the first step ACP S-acetyltransferase (ACOATA_c) was retained in MUST^U^. For the MUST^L^ set, downregulations of biomass-coupled reactions (i.e., 119) were excluded from further analysis since those perturbations reduce cellular growth for production gain. No flux pairs that considered sums and differences were identified by the analysis (MUST^UU^, MUST^LL^ and MUST^LU^ for overexpressed sum, downregulated sum, and overexpressed flux difference, respectively) [45]. This is due to the linearity of the acyl-CoA (from acetyl-CoA) and TAG synthesis pathways (i.e., sequential attachment of acyl-CoA to dihydroxyacetone backbone) and the absence of converging paths towards TAG. Overall, few perturbations (i.e., fourteen in MUST-single and none in MUST-pair) were suggested because under nitrogen limitation the reference wild-type fluxes already achieve a TAG production phenotype (though less than overproducing strain) and resemble the overproducing state. Surprisingly, we found that two key overproduction targets [70], (i) ATP citrate lyase and (ii) cytoplasmic malic enzyme (ME2_c), were not included in the MUST^U^ set. ATP citrate lyase is the key enzyme in producing cytoplasmic acetyl-CoA. However, acetyl-CoA synthetase can functionally replace ATP citrate lyase thus both reactions form a MUST^UU^ pair. However, since acetyl-CoA synthetase can participate in a high-flux thermodynamically infeasible cycle that transports acetyl-CoA to mitochondria, hydrolyzes it to acetate, exports to cytoplasm, and re-synthesizes acetyl-CoA, the ranges of the flux sum of the pair for wild-type and overproducing strain overlapped. This discovered cycle was subsequently removed from the model by turning off the ethanol-induced acetyl-CoA transport via carnitine shuttle [71] and ATP citrate lyase and acetyl-CoA synthetase were added to the MUST^UU^ set. Malic enzyme can participate in a transhydrogenase cycle involving malate dehydrogenase and pyruvate carboxylase and produce cytoplasmic NADPH for acyl-CoA synthesis. The model contains three mechanisms for cytoplasmic NADPH production hence the malic enzyme contribution can only be detected by looking at non-overlapped flux triplets. The other two mechanisms are oxidative pentose phosphate pathway (PPP) and glycerol dehydrogenase. All three NADPH production mechanisms were added to the MUST^U^ set and their overexpression levels were found by minimizing the reaction flux in the overproducing strain with the other two knocked out. Overall, there were ten candidates for overexpression (MUST^U^), three candidates for downregulation (MUST^L^), and no candidates for knockout (MUST^X^). All reactions in MUST sets are provided in Supplementary Materials 2. OptForce was used to search for combinations of candidates in MUST sets that can lead to enhanced TAG overproduction and recorded these combinations into the FORCE set (Table 4). Without the lipid degradation knockout, no combinations could be identified since OptForce min-max objective function identified the worst-case scenario of TAG synthesis-degradation cycle. beta-Oxidation was not originally placed in the MUST^X^ set since both wild-type and overproducing strains can run (to some extend) TAG and fatty acid synthesis-degradation cycles. The *PEX10* knockout [19, 68] was manually added along with the removal of fatty acid secretion since this phenotype was not observed in both wild type and engineered strains [11] and OptForce was rerun (see Table 4).

**Table 4.**
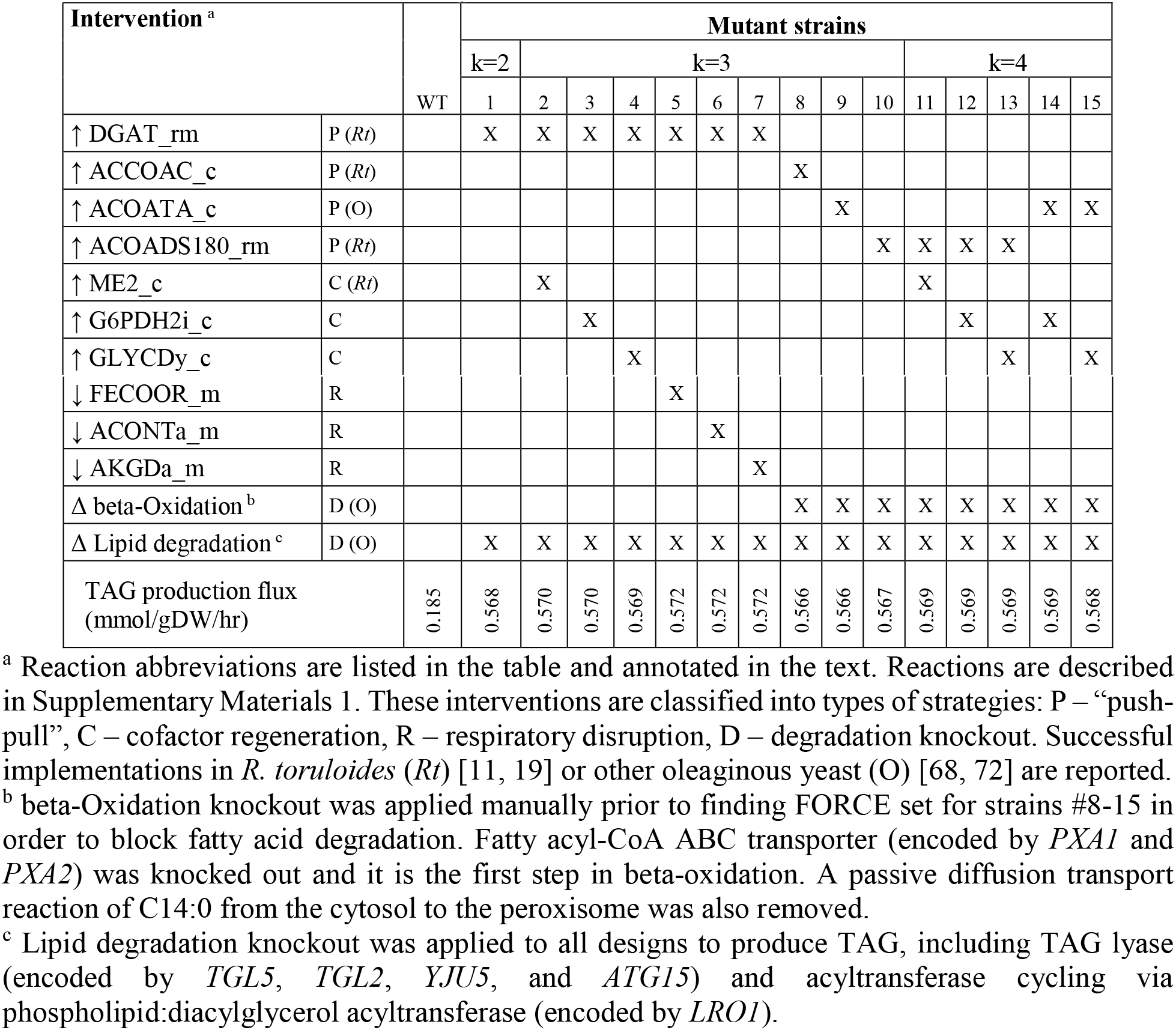
Combinations of genetic perturbations suggested by OptForce procedure for triacylglycerol production under nitrogen limitation.

OptForce identified a total of 15 sets of interventions (see Table 4). Several identified genetic interventions match successfully implemented strategies in *R. toruloides* (4 out of 12) [11, 19] and other oleaginous yeasts (3 out of 12) [68, 72]. Identified TAG overproducing strategies can be divided into three group: upregulation of precursor production and TAG production (“push-pull” strategies) [46], cofactor regeneration, and respiratory disruption (see Table 4). “Push-pull” interventions which directly increase the flux throughput present in all combinations. Cofactor regeneration and respiratory disruption interventions, which indirectly support TAG production, by themselves do not lead to robust TAG overproducing phenotypes. However, those interventions when applied in combination further increased TAG production for strains with “push-pull” interventions. When predicting the quantitative effectiveness of interventions, stoichiometric models cannot always capture the synergy of interventions strategies. For example, overexpression of diacylglycerol acyltransferase (DGAT_rm, *DGA1*) (Table 4) alone could achieve 89% of theoretical yield. However, both interventions, *DGA1* and *ACC1* (ACCOAC_c), were needed to derive a high-yield strain [11]. Overall, a maximum of two interventions per FORCE set (excluding manually applied degradation knockout interventions) were suggested by OptForce (see Table 4). Supplying NADPH for acyl-CoA synthesis via overexpressing cytoplasmic malic enzyme (*ME* or *rt4393*) has shown improvement as a single intervention but decreased lipid yield in the triple overexpression of *ACC1 DGA1 ME* [19]. Predictions with *iRhto*1108 showed a different trend where single interventions had little effect thus requiring double interventions following a “push-pull” strategy (e.g., *DGA1*) to improve yield. As discussed in Zhang et al., 2016 [11], a hypothesis for the counterintuitive behavior under *ME* overexpression is that increasing the flux through malic enzyme might disrupt cellular balance affect lipid biosynthesis and the transhydrogenase cycle. Knocking out fatty acid and lipid degradation pathways is another strategy that was proven effective in *Y. lipolytica* [68] and subsequently tested in *R. toruloides* [19]. This knockout was implemented by deleting gene *PEX10* required for peroxisome biogenesis. While the degradation pathway knockout is essential for robust TAG production using OptForce, *in vivo* implementation of Δ*PEX10* decreased both lipid fraction and biomass and lipid titer in *R. toruloides* [19]. Why peroxisome biogenesis contributes to cellular growth and lipid production is beyond the purview of a stoichiometric model such as *iRhto*1108.

We also compared the OptForce results with interventions that were implemented in other oleaginous organisms. As mentioned before, lipid and fatty acid degradation knockouts were effective in *Y. lipolytica* [68]. Overexpression of fatty acid synthase (with ACOATA_c being the first step) in the oleaginous fungus *Aspergillus oryzae* was found to increase fatty acid and TAG production by more than two-fold [72]. OptForce identified five new interventions that can increase TAG production when being applied in combination with a “push-pull” intervention. Two additional mechanisms for NADPH generation, glucose 6-phosphate dehydrogenase (G6PDH2i_c) in oxidative PPP or glycerol dehydrogenase (GLYCDy_c), are analogous to malic enzyme overexpression. Glycerol dehydrogenase upregulation is theoretically possible but introduces the toxic metabolite dihydroxyacetone [73]. On the other hand, glucose 6-phosphate dehydrogenase and more broadly oxidative PPP was found to be upregulated natively under nitrogen limitation in *Y. lipolytica* (two-fold flux increase in 13C-MFA study [69]) and *R. toruloides* (inferred by mutant phenotypes [19]). Thus, OptForce correctly identified the importance of oxidative PPP upregulation though a directed intervention may not be necessary. Finally, downregulation perturbations were suggested for aconitase (ACONTa_m), oxoglutarate dehydrogenase (AKGDa_m), or ferrocytochrome-c:oxygen oxidoreductase (FECOOR_m). These downregulations decrease cellular respiration, repress growth, and indirectly allow more carbon to be used in TAG production. Growth repression can also be achieved using nutrient limitation and culture optimization [68] while allowing the cell to maintain its growth robustness trait. Overall, *iRhto*1108-driven strain redesign using OptForce identifies many “push-pull” strategies. Not all strategies are in agreement with the experimental findings but upon careful interpretation of the desired metabolic redirections, alternative ways of achieving the same goal can be designed that bypass the specified interventions.

## Conclusions

In this work, we collect and organize functional genomics data [13] and prior knowledge into the genome-scale metabolic model *iRhto*1108. Essential cellular metabolism and growth capability of the model was validated extensively with experimental results, including gene essentiality [13] and growth data. *iRhto*1108 was also able to recapitulate experimentally-observed lipid accumulation phenotypes [15–17]. We showed that *iRhto*1108 can comprehensively capture *R. toruloides*’s metabolism and provide meaningful predictions that were validated with experimental data including suggestion of genetic perturbations leading to triacylglycerol overproducing strains. We envision that in the future *iRhto*1108 will aid in exploring the metabolic potential of *R. toruloides*, following in the footsteps of the model organisms *Saccharomyces cerevisiae* [74].

Despite careful curation, a large number of blocked reactions (i.e., 677 out of 2,203) remained in the model spanning multiple pathways. Most of them are transport reactions (i.e., 194 reactions) connecting the network. The rest participate in secondary metabolism and degradation of amino acid, fatty acid, and lipid. We chose to keep them in the hope that they would aid in gap filling attempts in the future. Stoichiometric models can capture all known interconversion routes from substrates to biomass components and products and globally balance cofactor needs. However, they inherently cannot mechanistically link enzyme levels with metabolite concentrations and metabolic fluxes. To this end, kinetic models offer a promising formalism for integrating such heterogeneous datasets [75]. Efforts towards this direction will require 13C-derived information on internal fluxes under a variety of genetic and environmental perturbations along with secreted products and biomass yield [76]. To this end, atom mapping models for all reactions in the *R. toruloides* model [77] will have to be constructed and robust methodologies for flux elucidation and kinetic model parameterization will have to be developed accounting for the multi-compartment nature of metabolism.

## Methods

### Draft reconstruction from existing fungal genome-scale reconstruction and model refinements

In general, the workflow used in this study followed an established protocol described in Mueller et al., 2013 [78] for generating a metabolic model utilizing a previously built metabolic model for a closely related organism. This protocol provides a priority structure for assigning functions to genes with multiple annotations. The most recent genome sequence and gene annotations of *R. toruloides* was used for this reconstruction [13]. The unannotated sequence of mitochondrial genome of *R. toruloides* was annotated using RAST [79] and MAP [80] on the KBase platform [42] (see Supplementary Materials 1). An initial draft reconstruction was assembled by mapping genes and reactions from the *S. cerevisiae* genome-scale model yeast 7.6 [40] with updated information from Chowdhury et. al., 2015 [81]. Briefly, first, homologous genes were determined by a bidirectional protein BLAST procedure [78] with an e-value cutoff of 10^−5^. The Boolean logic given by each gene-protein-reaction association (GPR) in yeast 7.6 was then evaluated using these bidirectional hits. A reaction was next added to the draft model only if its GPR satisfied can be satisfied with the present gene homologs necessary for a functional protein. This draft reconstruction was further extended with KBase’s “build fungal model” application [42] which extracts homologous genes and associated reactions from a library of fungal genome-scale models using similar homologous genes identification schematics. We prioritized building the initial scaffold using yeast 7.6 model rather than KBase because the biochemical information in yeast 7.6 was experimentally verified whenever possible. Next, additional reactions and GPRs were manually added using the annotated genome and validated with NCBI’s Conserved Domain Database [49]. Missing assignments of reaction compartments were resolved using the protein subcellular localization prediction software DeepLoc [82]. Adjustments made to reactions reversibility and activation in the default model are commented in Supplementary Materials 1 whereas other adjustments made specifically to model simulations are stated in the main text.

In addition to the biologically relevant additions and curations made to *iRhto*1108 (see Results and Methods), additional validations and refinements were performed to improve the model quality. Specifically, the GPRs of *S. cerevisiae* yeast 7.6 reactions that were recently updated in the yeast model repository (https://github.com/SysBioChalmers/yeast-GEM, version 8.3.3) were evaluated and modifications were made to the GPRs of 22 reactions. Furthermore, we ensured that every reaction is mass and charge balanced and as a result we updated 663 metabolite formulae (i.e., 33.5%) and 94 reaction stoichiometries (i.e., 4.4%) using standardized metabolite formulae from MetaCyc [83] and ModelSEED [84] databases. Database verification for metabolite formulae was at 77.8% coverage and we manually assigned the formulae assignments for the remainders to ensure all reactions were mass and charge balance (excluding pseudo and exchange reactions). Further model curation involved identifying and fixing thermodynamically infeasible cycles. For instance, cycles that allowed the unbounded production of ATP were eliminated by blocking the reverse direction of the ATP hydrolysis reactions [85]. *iRhto*1108 model structure was checked using the memote test suite [54] with the model annotations standardized to the MIRIAM namespace [86], which is used by memote. The final version of the model passed all memote tests.

### Generation of biomass reactions

Both experimental data from literature and those generated in this study (see Methods, Determination of biomass composition), alongside the original yeast 7.6 biomass reaction, were used to determine the metabolite coefficients in the biomass objective functions for carbon and nitrogen limitation conditions (Supplementary Materials 1). The macromolecular composition was measured for *R. toruloides* (see Methods, Determination of biomass composition). Other experimentally determined biomass specifications for *R. toruloides* were also incorporated, including genome GC content [58], lipid composition in nitrogen limitation conditions (i.e., natural, phospho, and glycolipids composition) [15], relative abundances of RNA nucleotide (this study), and relative abundance of acyl groups and free fatty acids in lipid (this study). Experimentally determined specifications taken from *S. cerevisiae* used in this model were amino acid, inorganic compound (phosphate, sulphate, and metal ions), and cell wall compositions [87], as corresponding data for *R. toruloides* was not available. Additional data adopted from the yeast 7.6 model were lipid subspecies composition (e.g., phosphatidylinositol, phosphatidylcholine, phosphatidylethanolamine, and phosphatidylserine composition). These data from experiments on *S. cerevisiae* and yeast 7.6 were deemed acceptable as *R. toruloides* and *S. cerevisiae* are closely related. The list of biomass constituents was reviewed and validated with relevant literature and experimental gene essentiality results (see Results). Without the measurement of the soluble metabolite pool, the coefficients of twelve cofactors and prosthetic groups were set to a small number of 10^−4^ so as to impose a biosynthesis requirement on the *in silico* model, resulting in 0.06% of the total biomass by weight. A similar measure was also adopted for the biomass reactions in other models such as those for *S. cerevisiae* models, including yeast 7.6 [62]. Metabolite coefficients associated with growth-associated ATP maintenance were also updated (see Methods, Determination of ATP maintenance requirements). Calculations and detailed listings of metabolites and coefficients in the biomass reaction are provided in the Supplementary Materials 1.

A total of 68 metabolites were included in the biomass component list for the *R. toruloides* model. The veracity of these inclusions was ascertained using data from single-gene knockout essentiality experiments in *R. toruloides* [13] and *S. cerevisiae* (see Supplementary Materials 2). Overlaps and differences between yeast 7.6 and *iRhto*1108’s list of biomass constituents are summarized in Table 1. Among the differences were eight nucleotide monophosphates which were replaced in *iRhto*1108 by the corresponding nucleotide triphosphates and pyrophosphate in order to directly account for DNA and RNA polymerization. The generic mannan metabolite was likewise substituted with N-glycan, O-glycan, and the glycosylphosphatidylinositol anchor [63, 88]. Similarly, the “generic” free fatty acid designation in yeast 7.6 was replaced with seven distinct free fatty acid compounds found with abundances of >1% by weight in the measurement of saponified fatty acids using LC-MS (see Methods, Experimental determination of biomass composition). In addition to the substitutions detailed above, nine cofactors and prosthetic groups suggested initially by Xavier et al., 2017 [44] were added to the list of biomass constituents so as to improve *iRhto*1108’s gene essentiality predictions. Seven new inorganic ions were also included in the biomass reaction following the measurements for *S. cerevisiae* by Lange and Heijnen, 2001 as these ions are known to be essential (Supplementary Materials 2). Although the biomass reaction in *iRhto*1108 is organism-specific, the list of biomass constituents is not unique to *iRhto*1108 and is applicable for the models of *S. cerevisiae* and possibly other closely related species such as *Y. lipolytica*. All newly added constituents in *iRhto*1108 were found to be essential for *S. cerevisiae* growth (see Supplementary Materials 2).

### Modeling simulation

Flux balance analysis (FBA) was used throughout the process for model validation and prediction [89]. Growth phenotypes were obtained using FBA with the objective of maximizing the biomass reaction (*v_biom_*) whose flux is equivalent to the growth rate. In general, the substrate uptake rates such as glucose (*v_glc_*) were set to the experimentally determined values if available. Otherwise, carbon substrate (e.g., glucose, xylose, or glycerol) uptake rate for a simulation was set to 5 mmol gDW^−1^ hr^−1^ which was close to the highest physiological glucose uptake rate found by Wang et al., 2018 [17]. For examining the model’s ability to utilize amino acid as nitrogen source, a specific amino acid uptake rate was set to 0.25 mmol gDW^−1^ hr^−1^ (i.e., 5% of default substrate uptake rate of 5 mmol gDW^−1^ hr^−1^). All simulations were performed using the carbon limitation condition model *iRhto*1108C unless *iRhto*1108N was specified to be used (i.e., for the nitrogen limitation condition).

For gene essentiality and mutant auxotrophy predictions in rich media [13], supplementary compound uptake rates were set to 0.25 mmol gDW^−1^ hr^−1^ (i.e., 5% of default substrate uptake rate of 5 mmol gDW^−1^ hr^−1^). Rich media components were described in Coradetti et al., 2018 [13] and are listed in Supplementary Materials 1. The undefined composition of yeast extract in Yeast-Peptone-Dextrose media was assumed to be that of YNB media plus 20 amino acids and d-glucose. The supplementary nutrients present in YNB included thiamine, riboflavin, nicotinate, pyridoxin, folate, (R)-pantothenate, 4-aminobenzoate, and myo-inositol. Oxygen and ammonium uptake rates were unconstrained in all simulations. Gene knockout was translated to the corresponding reaction(s) knockout by examining the Boolean gene-protein-reaction rules. A reaction was knocked out in the model by setting the corresponding upper and lower flux bounds to zero. A gene was determined to be essential if the knockout mutant’s maximal growth rate calculated by FBA was less than 0.0001 hr^−1^. The criteria for experimentally determined gene essentiality are described in Coradetti et al., 2018 [13]. The calculations were performed using the COBRApy package (version 0.13.4) [90].

### Determination of ATP maintenance requirements

Non-growth (NGAM) and growth associated ATP maintenance (GAM) values were determined using continuous chemostat data from Shen et al., 2013 [24]. A functional draft model utilizing a biomass reaction without a GAM demand was used to determine the biomass synthesis requirement excluding ATP maintenance. To calculate the ATP maintenance requirement per experimental data point, glucose uptake rate (*v_glc_*) and growth rate (*v_biom_*) were set to the experimentally determined values. Next, ATP maintenance requirement was given by the ATP hydrolysis rate (*v_atpm_*) which is the maximal through the following reaction: ATP + H_2_O → ADP + H^+^ + HPO_4_^2−^. An NGAM value of 1.01 mmol ATP gDW^−1^ hr^−1^ was found for no growth at *v_glc_* of 0.032 mmol gDW^−1^ hr^−1^, reported in Shen et al., 2013 [24]. GAM value was the slope of the line (found using linear regression) through all the maximal ATP maintenance rates constrained by the experimental *v_glc_*, *v_biom_*, and NGAM (by setting the intercept to the NGAM value). For nitrogen limitation condition, because the lipid composition of the biomass varied with the growth rate [24], per chemostats data point, the coefficients of the biomass reaction were adjusted in order to account for the compositional change. Specifically, lipid composition was adjusted to the experimentally determined value and the other macromolecular compositions (such as protein, carbohydrate, RNA, and DNA) were adjusted while maintaining the original relative levels. For carbon limitation conditions, the compositional profile remained relatively constant across the growth rates [15, 24] and thus no adjustments were made to the biomass reaction whilst determining the GAM value. Details of these simulations are provided in the Supplementary Materials 2.

### Phenotype phase plane and gene-flux correlation analysis

Phenotype phase planes [66] were used to calculate the maximal triacylglycerol yield under growth optimization priority under nutrient scarcity. Minimal oxygen, ammonium, phosphate, and sulphate uptake rates necessary for growth were found by minimizing the respective uptake rate subjected to maximal growth yield. The uptake rates were 12.7, 2.4, 0.20, and 0.03 mmol gDW^−1^ hr^−1^ for oxygen, ammonium, phosphate, and sulphate, respectively. Oxygen and ammonium (or phosphate or sulphate) uptake rates’ ranges were then discretized to 30 points between zero to the calculated minimal uptake value at maximal growth rate on the phenotype phase plane. For every point on the plane, a two-step procedure was applied. First, the growth rate was maximized subject to the limitation in oxygen and ammonium (or phosphate or sulphate) uptake rates. Second, triacylglycerol production was maximized subject to not only nutrient limitations but also first-priority growth optimization by constraining the biomass production to be at least the maximal amount determined in the previous step. These model-predicted phenotypes were used to construct the contour plots shown in Figure 2. TAG’s molecular weight of 882.40 mg mmol^−1^ derived from lipid’s acyl group composition was used in calculating TAG yield (unit of g TAG / g Glucose). Next, to compare the *in silico* flux redistribution in *iRhto*1108 (constructed for strain IFO0880) to the experimentally-observed differential gene expression in strain NP11 [3], we first established the mapping between NP11 genes and model reactions using bidirectional BLAST hit [78]. Once a mapping was established, the fold-change of the *in silico* metabolic flux can be calculated.

### Identification of genetic perturbation for triacylglycerol overproduction

The OptForce procedure was used to identify sets of genetic perturbations that once implemented result in strain with overproducing phenotypes, hereby called overproducing strains. Detailed formulations and explanations were documented in Ranganathan et al., 2010 [45] and additionally in Chowdhury et al., 2015 [91]. Throughout the simulation, diacylglycerol acyltransferase in lipid particle (DGAT_l) was turned off to allow flux to go through only endoplasmic reticulum version of the reaction (DGAT_rm). The first step in the OptForce procedure is to contrast flux ranges of a wild-type to those of an overproducing strain, one reaction at a time. Flux data obtained from 13C-MFA study of *Y. lipolytica* wild-type under nitrogen limitation [69] was used to constrain wild-type fluxes in this study. Specifically, 13C-MFA flux data for wild-type strain (among four sets of MFA fluxes) without separating cytoplasmic and mitochondrial malic enzyme fluxes were used because the study concluded that the compartmentalized malic enzyme fluxes were indistinguishable [69]. In the simulation, 13C-MFA flux range for malic enzyme were constrained on the sum of three reactions, namely cytoplasmic (ME2_c), mitochondrial with NAD+ (ME1_m), and mitochondrial with NADP+ (ME2_m) malic enzyme. The flux basis for this study was the glucose uptake of 10 mmol gDW^−1^ hr^−1^. We assess metabolic fluxes for both wild-type and overproducing strains under nitrogen limited conditions. Nitrogen availability level of 33% (0.7 mmol gDW^−1^ hr^−1^ in the basis of glucose uptake) was used in all OptForce simulations and was calculated by minimizing ammonium uptake rate for the wild-type strain with respect to reaction fluxes constrained by 13C-MFA data. Reference fluxes for overproducing strain were constrained by forced production of TAG to 90% of theoretical yield, growth yield required at 10% of wild-type, and aforementioned nitrogen limitation. In this work, flux variability analysis (FVA) [92] were used on *iRhto*1108 to generate the flux ranges for both wild-type and overproducing strains. Reactions with non-overlapping flux ranges were potential candidates for genetic perturbation. Reactions with FVA ranges being lower, higher, or equal zero (only) in overproducing strain compared to wild-type strain were put in MUST^L^, MUST^U^, or MUST^X^ set, respectively. Value ranges of flux sums (e.g., v_1_ + v_2_) and differences (e.g., v_1_ – v_2_) were also evaluated using FVA. Reaction pairs with higher values of flux sums, lower values of flux sums, and higher value of flux differences were put in MUST^UU^, MUST^LL^, and MUST^LU^ set, respectively. Reaction in the pair are put in the respective MUST^U^ or MUST^L^ depending on their membership in MUST paired set. The combinations of overexpression (MUST^U^), downregulations (MUST^L^), or knockout (MUST^X^) were determined using a bi-level mixed-integer linear programming optimization formulation and put in FORCE set [45]. Overexpression and downregulations were applied to the model by setting the reaction flux bound (lower and upper bound, respectively) to the level calculated in overproducing strain. Reaction knockout was applied by setting both the lower and upper bounds to zero. Because of the production minimization objective, OptForce could not suggest any overproducing strains due to lipid degradation pathways. To eliminate the TAG synthesis-degradation cycle that occurred under production minimization objective, TAG degradation and cycling pathways were knocked out in simulation, including neutral lipid lyase (6 reactions) and acyltransferase cycling via phospholipid:diacylglycerol acyltransferase (2 reactions). In addition, fatty acid secretions are not observed in *R. toruloides* in wild-type or engineered strains and thus were knocked out (12 reactions). For strains #8-15, the fatty acid degradation pathway was also blocked by knocking out the first step, fatty acyl-CoA ABC transporter to peroxisome (12 reactions) and tetradecanoate diffusion. The list of these knocked-out reactions are provided in the Supplementary Materials 2.

### Experimental determination of biomass composition

#### Strains and cultivation

Biomass composition was determined for *R. toruloides* IFO0880 cultivated in chemostat using two limiting nutrients (either glucose or nitrogen). An overnight stock was prepared in minimal medium that contains yeast nitrogen base without amino acid (YNB, Sigma Y0626) and 20g/L glucose. It was then inoculated into 250mL culture to grow in continuous mode in a 500-mL chemostat (Sixfors; Infors AG, Bottmingen, Switzerland). For the carbon limitation condition, YNB with 0.8g/L glucose was used. For the nitrogen limitation condition, 20g/L glucose and 0.05g/L (NH_4_)_2_SO_4_ were supplemented to YNB without ammonium sulfate (Difco, 291940). For both conditions, the culture was stirred at 400rpm for sufficient oxygen, and was kept at 0.1h^−1^ growth rate. After the culture had reached steady state (pH, oxygen, and cell density), it was harvested for the biomass measurement.

#### Biomass component analysis

DNA was measured using diphenylamine reagent. 7.5mL culture was pelleted and washed by 1mL cold 1 mM HClO_4_. Serial dilutions of 1mg/mL Calf thymus DNA (Sigma) were prepared for calibration. Samples were hydrolyzed in 500µL 1.6 M HClO_4_ for 30 min at 70°C, and then reacted with 1 ml diphenylamine reagent (0.5 g diphenylamine in 50 mL glacial acetate, 0.5 ml 98% H_2_SO_4_, and 0.125 ml 3.2 % acetaldehyde water solution) at 50°C for 3 hours. After centrifugation, the supernatant was taken for OD600 measurement.

RNA was measured by 260nm absorption. Basically, 2.5mL culture was pelleted and washed, and digested with 300µL 0.3M KOH at 37°C for 60 min. DNA and protein were then precipitated by 100 µL 3M HClO_4_. Supernatant was taken and precipitate was washed with 600µL 0.5M HClO_4_. Absorption of combined supernatant at 260nm, 900nm, and 977nm was measured. RNA concentration can be calculated as 5.6*(A260-A260blank)/(A977-A900) in µg/mL. RNA composition was calculated from the relative abundances of bases in RNA-Seq data. Extracted RNA was sequenced on a HiSeq2500 (Illumina). Fastq files were generated and demultiplexed with the bcl2fastq v1.8.4 Conversion Software Reads were trimmed by Trimmomatic [93] and analyzed using FastQC [94]. STAR version 2.5.4a [95] and featureCounts from the Subread package, version 1.5.2 [96] were used to map the reads to the *R. toruloides* strain IFO0880 reference genome [13] and obtain relative counts.

Protein was measured using the Biuret method. Briefly, 2.5mL culture was pelleted, washed and boiled in 100 µL 3M NaOH at 98°C for 5 min. After cool down, the mixture was reacted with 100 µL CuCO_4_ for 5 min at room temperature. After centrifugation, supernatant was taken for determination of 555nm absorption, which was calibrated by serial dilution of BSA solutions (Thermo).

Lipid was determined by measuring saponified fatty acids using LC-MS. Briefly, cell pellet from 2.5mL culture was extracted and saponified in 1mL 0.3M KOH-MeOH solution at 80°C for 60 min, then neutralized by 100 µL formic acid, and then extracted by 1mL hexane 20nM. 20nM isotope-labeled fatty acid standards (U-13C-C16:0, U-13C-C18:1, U-13C-C18:2; Cambridge Isotope) were added before saponification as internal standards. Extracted fatty acids were dried under N_2_ and redissolved in 200µL acetonitrile:methanol (1:1), and then analyzed by reversed-phase C8 column chromatography coupled to negative-ion mode, full-scan high-resolution LC-MS (Exactive, Thermo).

Carbohydrate was determined by hydrolyzing cell pellet from 2.5mL culture in 100µL 2M HCl at 80°C for 1 hr. 0.4mg U-^13^C-glucose was added as internal control before hydrolyzation. The lysate was neutralized by 100µL 2M NH_4_HCO_3_, diluted in 1.8mL 80% MeOH and centrifuged. The supernatant was taken and analyzed by negative-ion mode LC-MS equipped with hydrophilic interaction liquid chromatography (Q Exactive Plus, Thermo). Mass peaks equivalent to C_6_H_12_O_6_ were selected for quantification.

### Batch growth of R. toruloides IFO0880

#### Strain, media, and culture conditions

YPD medium (10 g/L yeast extract, 20 g/L peptone, and 20 g/L glucose) was used for routine growth of *R. toruloides* IFO0880. Growth rates of *R. toruloides* IFO0880 was tested in minimal medium (MM) using different C/N ratios. (20 g/L d-glucose, 1.7 g/L yeast nitrogen base without amino acids and ammonium sulfate, 0.4 - 7.1 g/L NH_4_Cl, C/N = 5:1 – 90:1, 25 mM Na_2_HPO_4_, 150 mM KH_2_PO_4_, pH 5.6). Stationary phase *R. toruloides* IFO0880 seed cultures were obtained by inoculating single colonies from a YPD agar plate into 25 mL YPD liquid medium in 125 mL baffled flask. For growth, the seed cultures were then used to inoculate into 50 mL minimal medium in 250 mL baffled flask with a starting OD600 of 1. The cells were then grown at 30℃ and 250 rpm. All experimental conditions were performed with four replicates. Growth viability on amino acids was tested in minimal media with different amino acids as nitrogen source (11.9 g/L alanine, 14 g/L serine, 5.8 g/L arginine, 9.7 g/L lysine, 15.3 g/L proline, 15.9 g/L threonine and 7.1 g/L NH_4_Cl as control). The growth viability of cellobiose, sucrose, and mannose were tested in modified media (70 g/L of carbon source, 10 g/L yeast extract, 1.7 g/L yeast nitrogen base without amino acids and ammonium sulfate, 8 g/L (NH_4_)_2_SO_4_, and 1 g/L MgSO_4_, pH 5.6).

#### Dry cell weight measurement

Cell growth was measured by the absorbance at 600 nm using cell density meter. Dry cell weights (DCW) were determined as follows. The 1-5 ml of culture samples were collected into pre-weighed tubes and centrifuged at 16,000 × g for 5 minutes. Supernatant was discarded, and pellets were then washed twice with 50 mM phosphate buffered saline. Washed pellets were dried till constant weight at 65℃ for 24 to 48 h. The tubes where then weighed.

## Declarations

### Availability of data and material

The models are provided in the Supplementary Materials 3 in MS-Excel spreadsheets, JSON, and SBML (version 3, level 1). A visual metabolic network map reconstructed using the Escher toolbox [97] is also available. The formulation of biomass reaction is available in Supplementary Materials 1. In addition, gene essentiality predictions, phenotype predictions, and OptForce strain design suggestions are all provided in the Supplementary Materials 2. The models, visual map, experimental data, and relevant prediction results were also uploaded to a memote-created repository (https://github.com/maranasgroup/iRhto_memote) [54]. The model was curated to address reconstruction issues found by memote test suite and to follow memote standardization [54]. All flux balance analysis simulations were performed using the COBRApy package [90]. OptForce [45] was performed using GAMS (version 24.8.5, GAMS Development Corporation) using IBM ILOG CPLEX solver (version 12.7.1.0) on the high-performance computing resource cluster of Pennsylvania State University’s Institute for CyberScience Advanced CyberInfrastructure (ICS-ACI).

### Authors’ contributions

Conceived the study: HZ CDM. Supervised study: CDM. Reconstructed the model: HVD SHJC. Measured biomass composition: YS TX JDR. Measured RNA base abundances and performed growth experiments: AD SSJ CVR. Performed model simulations: HVD. Analyzed the results: HVD PFS CDM. Wrote the paper: HVD PFS CDM. All authors read and approved the final manuscript.

## Acknowledgements

We would like to thank Lin Wang (Pennsylvania State University) for his advice on OptForce simulation, Debolina Sarkar (Pennsylvania State University) for her critical review of the manuscript, and Jose Juan Almagro Armenteros (Technical University of Denmark) for running DeepLoc to predict protein subcellular localization.

## Ethics approval and consent to participate

Not applicable.

## Consent for publication

Not applicable.

## References

1. Starmer WT, Fell JW, Catranis CM, Aberdeen V, Ma L-J, Zhou S, et al. Yeasts in the Genus Rhodotorula Recovered from the Greenland Ice Sheet. In: Rogers SO, Castello JD, editors. Life in Ancient Ice. Princeton University Press; 2005. p. 181–95. http://www.jstor.org/stable/j.ctt1dr350p.18.

2. Garay LA, Sitepu IR, Cajka T, Chandra I, Shi S, Lin T, et al. Eighteen new oleaginous yeast species. J Ind Microbiol Biotechnol. 2016;43:887–900. doi:10.1007/s10295-016-1765-3.

3. Zhu Z, Zhang S, Liu H, Shen H, Lin X, Yang F, et al. A multi-omic map of the lipid-producing yeast Rhodosporidium toruloides. Nat Commun. 2012;3:1112. doi:10.1038/ncomms2112.

4. Buzzini P, Innocenti M, Turchetti B, Libkind D, van Broock M, Mulinacci N. Carotenoid profiles of yeasts belonging to the genera Rhodotorula, Rhodosporidium, Sporobolomyces, and Sporidiobolus. Can J Microbiol. 2007;53:1024–31. doi:10.1139/W07-068.

5. Beopoulos A, Nicaud J-M, Gaillardin C. An overview of lipid metabolism in yeasts and its impact on biotechnological processes. Appl Microbiol Biotechnol. 2011;90:1193–206. doi:10.1007/s00253-011-3212-8.

6. Xue S-J, Chi Z-M, Zhang Y, Li Y-F, Liu G-L, Jiang H, et al. Fatty acids from oleaginous yeasts and yeast-like fungi and their potential applications. Crit Rev Biotechnol. 2018;38:1–12. doi:10.1080/07388551.2018.1428167.

7. Hu C, Zhao X, Zhao J, Wu S, Zhao ZK. Effects of biomass hydrolysis by-products on oleaginous yeast Rhodosporidium toruloides. Bioresour Technol. 2009;100:4843–7. doi:10.1016/J.BIORTECH.2009.04.041.

8. Li Y, Zhao Z (Kent), Bai F. High-density cultivation of oleaginous yeast Rhodosporidium toruloides Y4 in fed-batch culture. Enzyme Microb Technol. 2007;41:312–7. doi:10.1016/J.ENZMICTEC.2007.02.008.

9. Wiebe MG, Koivuranta K, Penttilä M, Ruohonen L. Lipid production in batch and fed-batch cultures of Rhodosporidium toruloides from 5 and 6 carbon carbohydrates. BMC Biotechnol. 2012;12:26. doi:10.1186/1472-6750-12-26.

10. Castañeda MT, Nuñez S, Garelli F, Voget C, De Battista H. Comprehensive analysis of a metabolic model for lipid production in Rhodosporidium toruloides. J Biotechnol. 2018;280:11–8. doi:10.1016/J.JBIOTEC.2018.05.010.

11. Zhang S, Skerker JM, Rutter CD, Maurer MJ, Arkin AP, Rao C V. Engineering *Rhodosporidium toruloides* for increased lipid production. Biotechnol Bioeng. 2016;113:1056–66. doi:10.1002/bit.25864.

12. Park Y-K, Nicaud J-M, Ledesma-Amaro R. The Engineering Potential of Rhodosporidium toruloides as a Workhorse for Biotechnological Applications. Trends Biotechnol. 2018;36:304–17. doi:10.1016/J.TIBTECH.2017.10.013.

13. Coradetti ST, Pinel D, Geiselman GM, Ito M, Mondo SJ, Reilly MC, et al. Functional genomics of lipid metabolism in the oleaginous yeast Rhodosporidium toruloides. Elife. 2018;7:e32110. doi:10.7554/eLife.32110.

14. Tchakouteu SS, Kopsahelis N, Chatzifragkou A, Kalantzi O, Stoforos NG, Koutinas AA, et al. *Rhodosporidium toruloides* cultivated in NaCl-enriched glucose-based media: Adaptation dynamics and lipid production. Eng Life Sci. 2017;17:237–48. doi:10.1002/elsc.201500125.

15. Shen H, Zhang X, Gong Z, Wang Y, Yu X, Yang X, et al. Compositional profiles of Rhodosporidium toruloides cells under nutrient limitation. Appl Microbiol Biotechnol. 2017;101:3801–9. doi:10.1007/s00253-017-8157-0.

16. Wu S, Zhao X, Shen H, Wang Q, Zhao ZK. Microbial lipid production by Rhodosporidium toruloides under sulfate-limited conditions. Bioresour Technol. 2011;102:1803–7. doi:10.1016/J.BIORTECH.2010.09.033.

17. Wang Y, Zhang S, Zhu Z, Shen H, Lin X, Jin X, et al. Systems analysis of phosphate-limitation-induced lipid accumulation by the oleaginous yeast Rhodosporidium toruloides. Biotechnol Biofuels. 2018;11:148. doi:10.1186/s13068-018-1134-8.

18. Bommareddy RR, Sabra W, Maheshwari G, Zeng A-P. Metabolic network analysis and experimental study of lipid production in Rhodosporidium toruloides grown on single and mixed substrates. Microb Cell Fact. 2015;14:36. doi:10.1186/s12934-015-0217-5.

19. Zhang S, Ito M, Skerker JM, Arkin AP, Rao C V. Metabolic engineering of the oleaginous yeast Rhodosporidium toruloides IFO0880 for lipid overproduction during high-density fermentation. Appl Microbiol Biotechnol. 2016;100:9393–405. doi:10.1007/s00253-016-7815-y.

20. Adrio JL. Oleaginous yeasts: Promising platforms for the production of oleochemicals and biofuels. Biotechnol Bioeng. 2017;114:1915–20. doi:10.1002/bit.26337.

21. Yu A-Q, Pratomo Juwono NK, Leong SSJ, Chang MW. Production of Fatty Acid-Derived Valuable Chemicals in Synthetic Microbes. Front Bioeng Biotechnol. 2014;2:78. doi:10.3389/fbioe.2014.00078.

22. Lee JJ., Chen L, Cao B, Chen WN. Engineering Rhodosporidium toruloides with a membrane transporter facilitates production and separation of carotenoids and lipids in a bi-phasic culture. Appl Microbiol Biotechnol. 2016;100:869–77. doi:10.1007/s00253-015-7102-3.

23. Jagtap SS, Rao C V. Production of d-arabitol from d-xylose by the oleaginous yeast Rhodosporidium toruloides IFO0880. Appl Microbiol Biotechnol. 2018;102:143–51. doi:10.1007/s00253-017-8581-1.

24. Shen H, Gong Z, Yang X, Jin G, Bai F, Zhao ZK. Kinetics of continuous cultivation of the oleaginous yeast Rhodosporidium toruloides. J Biotechnol. 2013;168:85–9. doi:10.1016/J.JBIOTEC.2013.08.010.

25. Thiele I, Palsson B. A protocol for generating a high quality genome-scale metabolic reconstruction. Nat Protoc. 2010;5:93–121.

26. O’Brien EJ, Monk JM, Palsson BO. Using genome-scale models to predict biological capabilities. Cell. 2015;161:971–87. doi:10.1016/j.cell.2015.05.019.

27. Maia P, Rocha M, Rocha I. *In silico* constraint-based strain optimization methods: The quest for optimal cell factories. Microbiol Mol Biol Rev. 2016;80:45–67. doi:10.1128/MMBR.00014-15.

28. Lee SY, Kim HU. Systems strategies for developing industrial microbial strains. Nat Biotechnol. 2015;33:1061–72.

29. Feist AM, Zielinski DC, Orth JD, Schellenberger J, Markus J, Palsson BØ. Model-driven evalution of the production potential for growth coupled products of Escherichia coli. Metab Eng. 2010;12:173–86.

30. Srinivasan S, Cluett WR, Mahadevan R. Constructing kinetic models of metabolism at genome-scales: A review. Biotechnol J. 2015;10:1345–59. doi:10.1002/biot.201400522.

31. King ZA, Lloyd CJ, Feist AM, Palsson BO. Next-generation genome-scale models for metabolic engineering. Curr Opin Biotechnol. 2015;35:23–9. doi:10.1016/J.COPBIO.2014.12.016.

32. Lopes H, Rocha I. Genome-scale modeling of yeast: chronology, applications and critical perspectives. FEMS Yeast Res. 2017;17. doi:10.1093/femsyr/fox050.

33. Österlund T, Nookaew I, Nielsen J. Fifteen years of large scale metabolic modeling of yeast: Developments and impacts. Biotechnol Adv. 2012;30:979–88. doi:10.1016/J.BIOTECHADV.2011.07.021.

34. Shi S, Zhao H. Metabolic Engineering of Oleaginous Yeasts for Production of Fuels and Chemicals. Front Microbiol. 2017;8:2185. doi:10.3389/fmicb.2017.02185.

35. Wei S, Jian X, Chen J, Zhang C, Hua Q. Reconstruction of genome-scale metabolic model of Yarrowia lipolytica and its application in overproduction of triacylglycerol. Bioresour Bioprocess. 2017;4:51. doi:10.1186/s40643-017-0180-6.

36. Kavšček M, Bhutada G, Madl T, Natter K. Optimization of lipid production with a genome-scale model of Yarrowia lipolytica. BMC Syst Biol. 2015;9:72. doi:10.1186/s12918-015-0217-4.

37. Kerkhoven EJ, Pomraning KR, Baker SE, Nielsen J. Regulation of amino-acid metabolism controls flux to lipid accumulation in Yarrowia lipolytica. NPJ Syst Biol Appl. 2016;2:16005. doi:10.1038/npjsba.2016.5.

38. Tiukova IA, Prigent S, Sandgren M, Nielsen J, Kerkhoven EJ. Genome-scale model of Rhodotorula toruloides metabolism. bioRxiv. 2019;:528489. doi:10.1101/528489.

39. Monk J, Nogales J, Palsson BO. Optimizing genome-scale network reconstructions. Nat Biotechnol. 2014;32:447–52. doi:10.1038/nbt.2870.

40. Aung HW, Henry SA, Walker LP. Revising the representation of fatty acid, glycerolipid, and glycerophospholipid metabolism in the consensus model of yeast metabolism. Ind Biotechnol. 2013;9:215–28. doi:10.1089/ind.2013.0013.

41. Kanehisa M, Goto S. KEGG: Kyoto Encyclopedia of Genes and Genomes. Nucleic Acids Res. 2000;28:27–30. doi:10.1093/nar/28.1.27.

42. Arkin AP, Cottingham RW, Henry CS, Harris NL, Stevens RL, Maslov S, et al. KBase: The United States Department of Energy Systems Biology Knowledgebase. Nat Biotechnol. 2018;36:566–9. doi:10.1038/nbt.4163.

43. Kumar VS, Maranas CD. GrowMatch: An Automated Method for Reconciling In Silico/In Vivo Growth Predictions. PLoS Comput Biol. 2009;5:e1000308. doi:10.1371/journal.pcbi.1000308.

44. Xavier JC, Patil KR, Rocha I. Integration of Biomass Formulations of Genome-Scale Metabolic Models with Experimental Data Reveals Universally Essential Cofactors in Prokaryotes. Metab Eng. 2017;39:200–8. doi:10.1016/J.YMBEN.2016.12.002.

45. Ranganathan S, Suthers PF, Maranas CD. OptForce: An Optimization Procedure for Identifying All Genetic Manipulations Leading to Targeted Overproductions. PLoS Comput Biol. 2010;6:e1000744. doi:10.1371/journal.pcbi.1000744.

46. Tai M, Stephanopoulos G. Engineering the push and pull of lipid biosynthesis in oleaginous yeast Yarrowia lipolytica for biofuel production. Metab Eng. 2013;15:1–9. doi:10.1016/J.YMBEN.2012.08.007.

47. Kot AM, Błażejak S, Gientka I, Kieliszek M, Bryś J. Torulene and torularhodin: “new” fungal carotenoids for industry? Microb Cell Fact. 2018;17:49. doi:10.1186/s12934-018-0893-z.

48. Sánchez BJ, Li F, Kerkhoven EJ, Nielsen J. SLIMEr: probing flexibility of lipid metabolism in yeast with an improved constraint-based modeling framework. bioRxiv. 2018;:324863. doi:10.1101/324863.

49. Marchler-Bauer A, Derbyshire MK, Gonzales NR, Lu S, Chitsaz F, Geer LY, et al. CDD: NCBI’s conserved domain database. Nucleic Acids Res. 2015;43:D222–6. doi:10.1093/nar/gku1221.

50. Koonin E V, Fedorova ND, Jackson JD, Jacobs AR, Krylov DM, Makarova KS, et al. A comprehensive evolutionary classification of proteins encoded in complete eukaryotic genomes. Genome Biol. 2004;5:R7. doi:10.1186/gb-2004-5-2-r7.

51. Evans CT, Ratledge C. Possible regulatory roles of ATP:citrate lyase, malic enzyme, and AMP deaminase in lipid accumulation by *Rhodosporidium toruloides* CBS 14. Can J Microbiol. 1985;31:1000–5. doi:10.1139/m85-189.

52. Kerscher SJ. Diversity and origin of alternative NADH:ubiquinone oxidoreductases. Biochim Biophys Acta - Bioenerg. 2000;1459:274–83. doi:10.1016/S0005-2728(00)00162-6.

53. Hsieh H-C, Kuan I-C, Lee S-L, Tien G-Y, Wang Y-J, Yu C-Y. Stabilization of d-amino acid oxidase from Rhodosporidium toruloides by immobilization onto magnetic nanoparticles. Biotechnol Lett. 2009;31:557–63. doi:10.1007/s10529-008-9894-z.

54. Lieven C, Beber ME, Olivier BG, Bergmann FT, Ataman M, Babaei P, et al. Memote: A community-driven effort towards a standardized genome-scale metabolic model test suite. bioRxiv. 2018;:350991. doi:10.1101/350991.

55. Schellenberger J, Lewis NE, Palsson BØ. Elimination of Thermodynamically Infeasible Loops in Steady-State Metabolic Models. Biophys J. 2011;100:544–53. doi:10.1016/J.BPJ.2010.12.3707.

56. Yang Z, Huang J, Geng J, Nair U, Klionsky DJ. Atg22 Recycles Amino Acids to Link the Degradative and Recycling Functions of Autophagy. Mol Biol Cell. 2006;17:5094–104. doi:10.1091/mbc.e06-06-0479.

57. Lange HC, Heijnen JJ. Statistical reconciliation of the elemental and molecular biomass composition of *Saccharomyces cerevisiae*. Biotechnol Bioeng. 2001;75:334–44. http://www.ncbi.nlm.nih.gov/pubmed/11590606. Accessed 20 Nov 2018.

58. . Nakase T, Komagatai K. DNA base composition of some species of yeasts and yeast-like fungi. 1971. https://www.jstage.jst.go.jp/article/jgam1955/17/5/17_5_363/_pdf. Accessed 20 Nov 2018.

59. Chan SHJ, Cai J, Wang L, Simons-Senftle MN, Maranas CD. Standardizing biomass reactions and ensuring complete mass balance in genome-scale metabolic models. Bioinformatics. 2017;33:3603–9. doi:10.1093/bioinformatics/btx453.

60. Feist AM, Palsson BO. The biomass objective function. Curr Opin Microbiol. 2010;13:344–9. doi:10.1016/J.MIB.2010.03.003.

61. Feist AM, Henry CS, Reed JL, Krummenacker M, Joyce AR, Karp PD, et al. A genome-scale metabolic reconstruction for *Escherichia coli* K-12 MG1655 that accounts for 1260 ORFs and thermodynamic information. Mol Syst Biol. 2007;3:121.

62. Mo ML, Palsson BØ, Herrgård MJ. Connecting extracellular metabolomic measurements to intracellular flux states in yeast. BMC Syst Biol. 2009;3:37. doi:10.1186/1752-0509-3-37.

63. Orlean P. Architecture and Biosynthesis of the Saccharomyces cerevisiae Cell Wall. Genetics. 2012;192:775–818. doi:10.1534/genetics.112.144485.

64. Liu H, Zhao X, Wang F, Jiang X, Zhang S, Ye M, et al. The proteome analysis of oleaginous yeast Lipomyces starkeyi. FEMS Yeast Res. 2011;11:42–51. doi:10.1111/j.1567-1364.2010.00687.x.

65. Probst K V., Schulte LR, Durrett TP, Rezac ME, Vadlani P V. Oleaginous yeast: a value-added platform for renewable oils. Crit Rev Biotechnol. 2016;36:942–55. doi:10.3109/07388551.2015.1064855.

66. Edwards JS, Ramakrishna R, Palsson BO. Characterizing the metabolic phenotype: A phenotype phase plane analysis. Biotechnol Bioeng. 2002;77:27–36. doi:10.1002/bit.10047.

67. Chubukov V, Uhr M, Le Chat L, Kleijn RJ, Jules M, Link H, et al. Transcriptional regulation is insufficient to explain substrate-induced flux changes in Bacillus subtilis. Mol Syst Biol. 2013;9:709. doi:10.1038/msb.2013.66.

68. Blazeck J, Hill A, Liu L, Knight R, Miller J, Pan A, et al. Harnessing Yarrowia lipolytica lipogenesis to create a platform for lipid and biofuel production. Nat Commun. 2014;5:3131. doi:10.1038/ncomms4131.

69. Wasylenko TM, Ahn WS, Stephanopoulos G. The oxidative pentose phosphate pathway is the primary source of NADPH for lipid overproduction from glucose in Yarrowia lipolytica. Metab Eng. 2015;30:27–39. doi:10.1016/J.YMBEN.2015.02.007.

70. Liang M-H, Jiang J-G. Advancing oleaginous microorganisms to produce lipid via metabolic engineering technology. Prog Lipid Res. 2013;52:395–408. doi:10.1016/J.PLIPRES.2013.05.002.

71. Schmalix W, Bandlow W. The ethanol-inducible YAT1 gene from yeast encodes a presumptive mitochondrial outer carnitine acetyltransferase. J Biol Chem. 1993;268:27428–39. http://www.ncbi.nlm.nih.gov/pubmed/8262985. Accessed 29 Mar 2019.

72. Tamano K, Bruno KS, Karagiosis SA, Culley DE, Deng S, Collett JR, et al. Increased production of fatty acids and triglycerides in Aspergillus oryzae by enhancing expressions of fatty acid synthesis-related genes. Appl Microbiol Biotechnol. 2013;97:269–81. doi:10.1007/s00253-012-4193-y.

73. Molin M, Norbeck J, Blomberg A. Dihydroxyacetone kinases in Saccharomyces cerevisiae are involved in detoxification of dihydroxyacetone. J Biol Chem. 2003;278:1415–23. doi:10.1074/jbc.M203030200.

74. Borodina I, Nielsen J. Advances in metabolic engineering of yeast *Saccharomyces cerevisiae* for production of chemicals. Biotechnol J. 2014;9:609–20. doi:10.1002/biot.201300445.

75. Chowdhury A, Khodayari A, Maranas CD. Improving prediction fidelity of cellular metabolism with kinetic descriptions. Curr Opin Biotechnol. 2015;36:57–64. doi:10.1016/J.COPBIO.2015.08.011.

76. Khodayari A, Maranas CD. A genome-scale Escherichia coli kinetic metabolic model k-ecoli457 satisfying flux data for multiple mutant strains. Nat Commun. 2016;7:13806. doi:10.1038/ncomms13806.

77. Gopalakrishnan S, Maranas C, Gopalakrishnan S, Maranas CD. Achieving Metabolic Flux Analysis for S. cerevisiae at a Genome-Scale: Challenges, Requirements, and Considerations. Metabolites. 2015;5:521–35. doi:10.3390/metabo5030521.

78. Mueller TJ, Berla BM, Pakrasi HB, Maranas CD. Rapid construction of metabolic models for a family of Cyanobacteria using a multiple source annotation workflow. BMC Syst Biol. 2013;7:142. doi:10.1186/1752-0509-7-142.

79. Brettin T, Davis JJ, Disz T, Edwards RA, Gerdes S, Olsen GJ, et al. RASTtk: A modular and extensible implementation of the RAST algorithm for building custom annotation pipelines and annotating batches of genomes. Sci Rep. 2015;5:8365. doi:10.1038/srep08365.

80. Huntemann M, Ivanova NN, Mavromatis K, Tripp HJ, Paez-Espino D, Tennessen K, et al. The standard operating procedure of the DOE-JGI Metagenome Annotation Pipeline (MAP v.4). Stand Genomic Sci. 2016;11:17. doi:10.1186/s40793-016-0138-x.

81. Chowdhury R, Chowdhury A, Maranas C. Using Gene Essentiality and Synthetic Lethality Information to Correct Yeast and CHO Cell Genome-Scale Models. Metabolites. 2015;5:536–70. doi:10.3390/metabo5040536.

82. Almagro Armenteros JJ, Sønderby CK, Sønderby SK, Nielsen H, Winther O. DeepLoc: prediction of protein subcellular localization using deep learning. Bioinformatics. 2017;33:3387–95. doi:10.1093/bioinformatics/btx431.

83. Caspi R, Billington R, Fulcher CA, Keseler IM, Kothari A, Krummenacker M, et al. The MetaCyc database of metabolic pathways and enzymes. Nucleic Acids Res. 2018;46:D633–9. doi:10.1093/nar/gkx935.

84. Henry CS, DeJongh M, Best AA, Frybarger PM, Linsay B, Stevens RL. High-throughput generation, optimization and analysis of genome-scale metabolic models. Nat Biotechnol. 2010;28:977–82. doi:10.1038/nbt.1672.

85. Fritzemeier CJ, Hartleb D, Szappanos B, Papp B, Lercher MJ. Erroneous energy-generating cycles in published genome scale metabolic networks: Identification and removal. PLOS Comput Biol. 2017;13:e1005494. doi:10.1371/journal.pcbi.1005494.

86. Juty N, Le Novere N, Laibe C. Identifiers.org and MIRIAM Registry: community resources to provide persistent identification. Nucleic Acids Res. 2012;40:D580–6. doi:10.1093/nar/gkr1097.

87. Klis FM, de Koster CG, Brul S. Cell wall-related bionumbers and bioestimates of Saccharomyces cerevisiae and Candida albicans. Eukaryot Cell. 2014;13:2–9. doi:10.1128/EC.00250-13.

88. Goto M. Protein *O*-Glycosylation in Fungi: Diverse Structures and Multiple Functions. Biosci Biotechnol Biochem. 2007;71:1415–27. doi:10.1271/bbb.70080.

89. Orth JD, Thiele I, Palsson BØ. What is flux balance analysis? Nat Biotechnol. 2010;28:245–8. doi:10.1038/nbt.1614.

90. Ebrahim A, Lerman JA, Palsson BO, Hyduke DR. COBRApy: COnstraints-Based Reconstruction and Analysis for Python. BMC Syst Biol. 2013;7:74.

91. Chowdhury A, Zomorrodi AR, Maranas CD. Bilevel optimization techniques in computational strain design. Comput Chem Eng. 2015;72:363–72. doi:10.1016/J.COMPCHEMENG.2014.06.007.

92. Mahadevan R, Schilling CH. The effects of alternate optimal solutions in constraint-based genome-scale metabolic models. Metab Eng. 2003;5:264–76. doi:10.1016/J.YMBEN.2003.09.002.

93. Bolger AM, Lohse M, Usadel B. Trimmomatic: a flexible trimmer for Illumina sequence data. Bioinformatics. 2014;30:2114–20. doi:10.1093/bioinformatics/btu170.

94. Andrews S. FastQC: a quality control tool for high throughput sequence data. 2010. http://www.bioinformatics.babraham.ac.uk/projects/fastqc.

95. Dobin A, Davis CA, Schlesinger F, Drenkow J, Zaleski C, Jha S, et al. STAR: ultrafast universal RNA-seq aligner. Bioinformatics. 2013;29:15–21. doi:10.1093/bioinformatics/bts635.

96. Liao Y, Smyth GK, Shi W. featureCounts: an efficient general purpose program for assigning sequence reads to genomic features. Bioinformatics. 2014;30:923–30. doi:10.1093/bioinformatics/btt656.

97. King ZA, Dräger A, Ebrahim A, Sonnenschein N, Lewis NE, Palsson BO. Escher: A Web Application for Building, Sharing, and Embedding Data-Rich Visualizations of Biological Pathways. PLOS Comput Biol. 2015;11:e1004321. doi:10.1371/journal.pcbi.1004321.

